# Cell type-independent profiling of interactions between intracellular pathogens and the human phosphoproteome

**DOI:** 10.1101/2022.09.27.509702

**Authors:** Kyle Mohler, Jack Moen, Svetlana Rogulina, Jesse Rinehart

**Affiliations:** Department of Cellular & Molecular Physiology, Yale School of Medicine, New Haven, CT 06520, USA; Systems Biology Institute, Yale University, New Haven, CT 06516, USA.

**Keywords:** Host-Pathogen Interactome, Phosphoprotein, SARS-CoV-2

## Abstract

Interactions between proteins from intracellular pathogens and host proteins in an infected cell are often mediated by post-translational modifications encoded in the host proteome. Identifying protein modifications, such as phosphorylation, that dictate these interactions remains a defining challenge in unraveling the molecular mechanisms of pathogenesis. We have developed a platform in engineered bacteria that displays over 110,000 phosphorylated human proteins coupled to a fluorescent reporter system capable of identifying the host-pathogen interactome of phosphoproteins (H-PIP). This resource broadly enables cell-type independent interrogation and discovery of proteins from intracellular pathogens capable of binding phosphorylated human proteins. As an example of the H-PIP platform, we generated a unique, high-resolution SARS-CoV-2 interaction network which expanded our knowledge of viral protein function and identified understudied areas of host pathology.

## Introduction

Protein-protein interactions (PPIs) are essential components of the cellular environment, enabling protein complex assembly and facilitating the regulation of protein function. Protein interactions across the host proteome are finely tuned and tightly regulated both spatially and temporally by protein phosphorylation and other post-translational modifications (PTMs). Protein phosphorylation provides the cell with a means to rapidly alter cellular processes without requiring changes to proteome composition. This dynamically transient regulatory strategy positions protein phosphorylation as an efficient modulator of cellular function, but consequently confounds systems-wide studies of phosho-dependent PPIs in their native cellular environment (Humphrey et al., 2015; Olsen et al., 2006). At the fundamental level, the sensitivity and efficiency that define successful phosho-regulatory mechanisms creates an intrinsic susceptibility of the global regulatory network to disruption and exploitation by exogenous protein interactions, like those initiated by proteins from viruses and other intracellular pathogens (Suryawanshi et al., 2021).

Intracellular pathogens use relatively small genomes to produce functionally diverse proteins that act collectively to redirect host cellular processes, evade detection, and promote replication (Banerjee et al., 2020). Traditional approaches to identification of host-pathogen interactions effectively generate large PPI data sets through physical capture of stable host-pathogen protein complexes, but their comprehensiveness is technically limited by the abundance of interacting host proteins, availability of *in vitro* cell models, and stability of protein complexes. In addition to transient PPIs, these approaches fail to identify sequence-specific binding sites between interacting proteins and are consequently devoid of interactions facilitated by specific protein phosphorylation events.

The successful implementation of a phospho-dependent PPI analysis platform hinges on a reliable and robust method for the production of phosphoproteins. The standard methods of creating a phosphorylated protein either rely on the use of a secondary enzyme, e.g. protein kinases, or make use of orthogonal translation systems (OTSs). The use of protein kinases is practically limited to previously characterized kinase-substrate interactions, and technically limited by issues of protein expression, off-target activity, and scalability in recombinant systems. OTSs bypass many of these limitations by enabling the specific co-translational insertion of a phospho-amino acid during recombinant protein expression. The phosphoserine-OTS (pSerOTS) makes use of a phosphoseryl-tRNA synthetase (pSerRS) derived from *Methanococcus meriplaudis* to aminoacylate pSer onto a UAG-decoding suppressor tRNA^pSer^ modified from *Methanococcus janaschii* tRNA^Cys^. To facilitate delivery of pSer-tRNA^pSer^ to the ribosome, *E. coli* elongation factor Tu (EF-Tu) variants (EF-pSer) have been evolved to bind the larger, negatively charged pSer moiety (Figure 1A) (Park et al., 2011). While advances in OTS technology have expanded their utility in recombinant protein expression, their deployment in complex, high-throughput PPI investigation strategies has been limited by practical considerations and deleterious interactions with native host systems.

**Figure 1.**
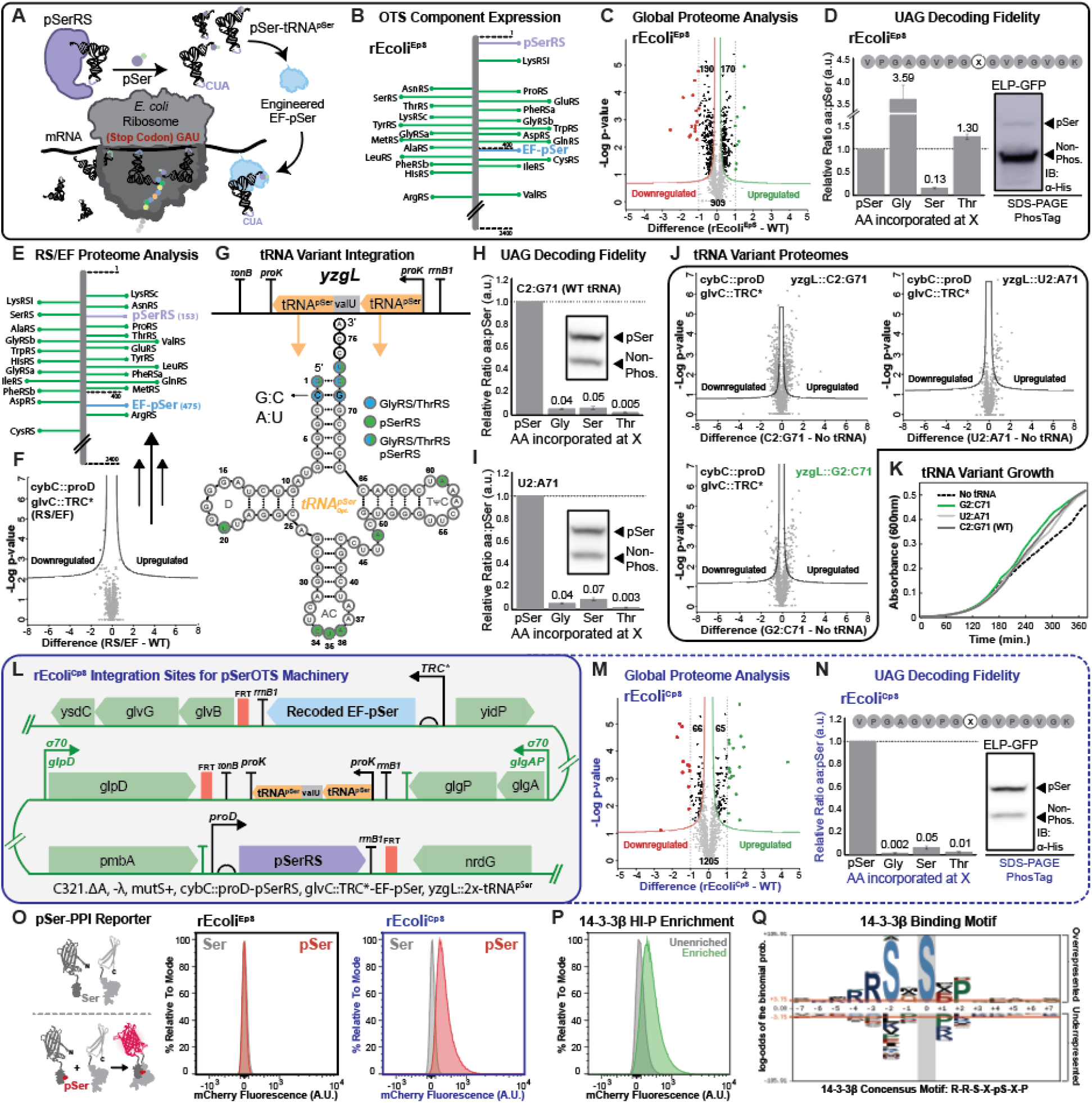
Stable integration of pSerOTS enables complex pSer-mediated PPI investigation. (**A**) Production of pSer-containing proteins using orthogonal phosphoseryl-tRNA synthetase (pSerRS) and tRNA pair. (**B**) Relative protein expression levels (rank ordered aaRSs from proteome, green) for pSerRS (purple) and EF-pSer (blue) in cells with episomal pSerOTS. (**C**) Volcano plot depicting changes in protein expression in rEcoli^EpS^ compared to cells without episomal pSerOTS (WT). Proteins above the curved lines represent significant expression changes with upregulated (Green) and downregulated (Red) proteins >1-fold change indicated; n = 3. (**D**) Mass spectrometry-based reporter for translation fidelity at TAG codons in rEcoli^EpS^. Bar graph displaying values for Gly, Ser, and Thr misincorporation relative to pSer (represented by the non-phospho band on the PhosTag immunoblot), n=3. (**E**) Relative protein expression levels (rank ordered aaRSs from proteome, green) for pSerRS (purple) and EF-pSer (blue) in cells with integrates pSerRS and EF-pSer (RS/EF). (**F**) Volcano plot depicting changes in protein expression in RS/EF cells compared to cells without integrated components (WT). Proteins above the curved lines represent significant expression changes; n = 3. (**G**) Schematic of tRNA integration cassette with tRNA primary sequence and secondary structure highlighting pSerRS (green) and GlyRS/ThrRS (blue) recognition elements and site targeted for mutation (C2:G71). (**H-I**) Mass spectrometry-based reporter for translation fidelity at TAG codons in cells with integrated tRNA variant cassettes. Bar graph displaying values for Gly, Ser, and Thr misincorporation relative to pSer (represented by the non-phospho band on the PhosTag immunoblot), n=3. (**J**) Volcano plot depicting changes in protein expression between integrated tRNA variant strains and cells without integrated components (WT). Proteins above the curved lines represent significant expression changes; n = 3. (**K**) Growth analysis of strains with integrated tRNA variant cassettes (green and gray lines) compared to WT tRNA (black line) and the RS/EF background strain (black dashed line). (**L**) Integration sites for pSerOTS components in rEcoli^CpS^. (**M**) Volcano plot depicting changes in protein expression in rEcoli^CpS^ compared to a strain without integrated components (WT). Proteins above the curved lines represent significant expression changes with upregulated (Green) and downregulated (Red) proteins >1-fold change indicated; n = 3. (**N**) Mass spectrometry-based reporter for translation fidelity at TAG codons in rEcoli^CpS^. Bar graph displaying values for Gly, Ser, and Thr misincorporation relative to pSer (represented by the non-phospho band on the PhosTag immunoblot), n=3. (**O**) Flow cytometry-based split-mCherry reporter assay for pSer incorporation for episomal pSerOTS (black) compared to the same reporter in rEcoli^CpS^ (blue). pSer incorporation facilitates PPI and reconstitutes mCherry fluorescence. (**P**) Flow cytometry analysis of population enrichment for pSer-mediated 14-3-3β interactions using H-PIP. (**Q**) Motif enrichment using pLogo analysis of phosphosites identified from enriched population in (F) match known consensus motif.

To enable the first comprehensive investigation pipeline for pSer-mediated pathogen-host PPIs, we engineered a strain of genomically recoded *E. coli* which can site-specifically incorporate pSer during protein translation via a chromosomally integrated and constitutively active pSerOTS. We then coupled our newly constructed strain with a modified version of our recently described PPI framework (Barber et al., 2018), which together establish a pipeline to facilitate the cell type-independent identification of phosphorylation-mediated host-pathogen PPIs with phosphosite resolution (Fig. 1A). To illustrate feasibility of our approach, we characterized pSer-mediated SARS-CoV-2 PPIs across the human proteome and generated a unique, high-resolution viral protein interaction network which expanded known viral protein function and our understanding of the mechanisms used by viral proteins to subvert or alter host cellular processes.

## Results

### A genetically modified *E. coli* strain enhances pSer-mediated PPI screening

The success of our approach was dependent on the modification and optimization of foundational phosphoprotein production technologies hindered by extraneous demands on native protein translation, enhanced metabolic burden, and conditional component orthogonality (Lee et al., 2013; Pirman et al., 2015; Rogerson et al., 2015; Zhu et al., 2019). Systematic evaluation of pSerOTS components in our lab identified pSerRS overexpression as the primary effector of host toxicity and decreases in OTS performance (Mohler et al., 2021). Steady-state levels of pSerRS were assessed by shotgun proteomics and ranked relative to homologous translational machinery (i.e. aaRSs) and the broader host proteome. From an episomal vector, pSerRS can become one of the most abundant (top ten) proteins in the cell, placing expression well above the level of native aminoacyl-tRNA synthetases (aaRS) (Figure 1B). When compared to the genomically recoded background strain (rEcoli) (Lajoie et al., 2013), the proteomes of cells harboring episomal pSerOTS (rEcoli^EpS^) were highly perturbed, with significantly altered expression of ∼400 proteins related to stress response pathway activation and metabolic reprogramming (Figure 1C).

In addition to increased translational demand, high-level aaRS expression has been shown to compromise aminoacyl-tRNA pool fidelity, thus decreasing translational accuracy and synergistically enhancing cellular stress (Kelly et al., 2019; Mohler et al., 2017b; Mohler et al., 2018; Sherman et al., 1992). To illustrate the impact of this cellular environment on OTS performance, we used an optimized mass spectrometry reporter for exact amino acid decoding (MS-READ) (Mohler et al., 2017a) to monitor OTS-mediated pSer incorporation and decoding fidelity in MS-READ protein expressed in rEcoli^EpS^ cells. We observed a strikingly low level of pSer incorporation relative to standard amino acids misincorporations (pSer(1):Gly(3.6):Thr(1.3) ratio), Figure 1D) at the same position, indicative of compromised OTS performance. Immunoblot analysis of the MS-READ protein following resolution by PhosTag SDS-PAGE (which allows physical separation of phosphoproteins from their non-phosphorylated counterpart) confirmed that the majority of MS-READ protein expressed was non-phosphorylated (Figure 1D). Closer inspection of OTS vectors isolated from rEcoli^EpS^ cell populations with compromised pSerOTS function revealed frequent inactivating transposon insertion events which improved OTS vector tolerance by ablating OTS function (**Figure S1. Related to** Figure 1). Based on these results, we concluded that OTS stability was consistently compromised by persistent negative selection pressure in rEcoli^EpS^ cells and that genomic integration of pSerOTS components would be an efficient means to control component expression and bypass the deleterious effects and intrinsic vulnerabilities of episomal OTS expression.

The foundation of the integrated pSerOTS strain is a genomically recoded strain of *E. coli* C321.ΔA (rEcoli) modified to replace every genomic occurrence of the amber (UAG) stop codon with the UAA stop codon (Isaacs et al., 2011; Lajoie et al., 2013). Together with the deletion of UAG-terminating release factor 1 (RF1), this strain opens the UAG codon to use by pSerOTS and reduces cellular stress caused by off-target nonsense suppression in non-recoded cells (Heinemann et al., 2012; Johnson et al., 2012; Johnson et al., 2011). To create a stable intracellular pool of pSer for aminoacylation, we deleted the gene encoding the *E .coli* Ser phosphatase, *serB*, which natively converts pSer to Ser during Ser biosynthesis (Steinfeld et al., 2014). Using this strain as the background, we genomically integrated individual pSerOTS component-promoter pairs to systematically tune component expression to previously determined levels which optimally balance OTS performance with cellular viability (Mohler et al., 2021).

pSerRS, EF-pSer, and tRNA^pSer^ comprise the essential components of the pSerOTS that must be integrated to facilitate function (Figure 1A). To minimize the impact of localized increases in transcriptional and translational demand at component integration sites and bypass expression variability associated with polycistronic operons, we distributed component integration across three separate pseudogene loci: *cybC*, *glvC*, and *yzgL* (Meyer et al., 2019; Trower, 1993). We aimed to install the pSerOTS as a natively active complement to the host translational machinery and selected four constitutively active transcriptional promoters with increasing strength (*glnSp\**, *OXB20*, *proDp*, *TRC*)* to drive steady-state expression of integrated OTS components across a wide proteomic range. Following genomic integration of promoter-pSerRS variant cassettes at the *cybC* locus, steady-state levels of pSerRS were assessed by shotgun proteomics and ranked in context of host proteome. As expected, pSerRS promoter variants enabled expression across a wide proteomic range with the lowest relative expression driven by *glnSp*-*pSerRS (∼1900) and the highest by *proDp*-pSerRS (∼190), mirroring known relative transcriptional activity (**Figure S2A-B. Related to** Figure 1) (Chatterjee et al., 2013). Both *TRC\**- and *proDp*-pSerRS fell within the relative range of endogenous aaRSs, however, the *proDp*-pSerRS strain was selected for continued engineering after comparative proteomic analysis revealed significant proteomic perturbation in *TRC\**-but not *proDp*-pSerRS strain backgrounds (**Figure S2C. Related to** Figure 1).

Integration and analysis of EF-pSer followed a similar workflow, but the *glnSp\** promoter variant was excluded due to low-level genomic expression. The primary genomic sequence of EF-pSer was recoded to prevent off-target integration at the native EF-Tu locus and directed towards integration at *glvC* (**Figure S2D. Related to** Figure 1). Proteomic analysis of integrated promoter-EF-pSer variant strains identified a high correlation between the steady-state EF-pSer expression profiles and those observed for pSerRS, however, *proDp*-EF-pSer integration dramatically altered host proteome composition compared to the intermediate expression *TRC\**-EF-pSer (∼640) variant strain selected for subsequent tRNA component integration (Figure 1F**; Figure S2E-G. Related to** Figure 1).

Two copies of tRNA^pSer^ were integrated at the *yzgL* locus in a transcriptionally insulated, optimized tRNA cassette with a native valU tRNA linker to enhance tRNA processing following transcription driven by a native proK tRNA promoter-terminator pair (Chatterjee et al., 2013). We previously observed that the orthogonality of tRNA expressed from an episomal OTS could be improved through base pair substitution of a primary host aaRS recognition element (C2:G71) in the tRNA^pSer^ acceptor stem (Mohler et al., 2021) (Figure 1G). We assessed the impact of unmodified (C2:G71) and two modified (A2:U71, G2:C71) tRNA variant cassette integrations on decoding fidelity using MS-READ and on the broader host proteome by comparative proteomics. All integrated tRNA variants showed dramatically improved decoding fidelity compared to the episomal pSerOTS (Figure 1G-I**; Figure S2I. Related to** Figure 1). When compared to the unmodified tRNA variant cassette (Figure 1H) the A2:U71 modified tRNA (Figure 1I) had similar decoding fidelity, but the G2:C71 tRNA variant cassette showed vastly improved fidelity (∼20-fold reduction in Gly misincorporation) (Figure 1N). Both modified tRNA variants reduced alterations to the host proteome caused by tRNA integration (Figure 1J), but the G2:C71 tRNA variant strain had enhanced growth properties when compared to the A2:U71 tRNA integrant strain (Figure 1K) and was selected as the final pSerOTS strain.

The new recoded *E. coli* strain with chromosomally integrated pSer machinery (rEcoli^CpS^) provides pSer as a 22nd amino acid available for translation at UAG codons (Figure 1L). Compared to episomal pSerOTS, rEcoli^CpS^ is stable, easy to use, and facilitates efficient production of pSer-containing proteins while reducing significant alterations to the host proteome by ∼3-fold (Figure 1C; Figure 1M). Additionally, the yield and purity of pSer-containing recombinant protein expressed in rEcoli^CpS^ cells is greatly improved compared to the same protein expressed in rEcoli^EpS^, with ∼1000-fold enhancement (Figure 1D; Figure 1N). Importantly, we found that rEcoli^CpS^, but not rEcoli^EpS^, was able to reliably facilitate pSer-mediated PPIs using a phospho-dependent split mCherry reporter system (Figure 1O) and recapitulate PPIs identified using sub-optimal episomal pSerOTS systems by robustly enriching PPIs of the known pSer-binding protein 14-3-3β and generating a canonical 14-3-3 binding motif (**Figure P-Q**) (Barber et al., 2018).

### pSer-mediated host-pathogen PPIs form functionally enriched networks

To enable discovery and deconvolution of organism-wide host-pathogen interactions, we coupled rEcoli^CpS^ cells to a high-throughput PPI detection platform capable of identifying phosphosite-specific protein interactions to establish the host-pathogen interactome of phosphoproteins (H-PIP) pipeline. As a demonstration of the H-PIP pipeline, we characterized the pSer-mediated protein interactions of fifteen SARS-CoV-2 proteins across the human proteome. The PPI detection platform is based on bimolecular fluorescence complementation using a split mCherry fluorescent protein reporter (Sawyer et al., 2014). The C-terminal half of a split mCherry was fused to an each of the fifteen individual viral proteins (or protein domains), forming the backbones of a PPI libraries. Each of the 110,139 annotated pSer phosphosite was then introduced into PPI each PPI vector as individual fusions with the N-terminal domain of the split mCherry. The fully assembled PPI libraries enabled the simultaneous pairwise analysis of each phosphosite with an individual SARS-CoV-2 protein. Flow cytometry was used to analyze individual cells in the population and identify productive interactions by measuring shifts in fluorescence intensity that result from PPI-mediated reconstitution of the fluorescent protein. Fluorescent cells containing productive interactions were separated from non-productive cells in the population, enriched over several rounds, and identified by next generation sequencing of phosphosites in the enriched population (Figure 2A).

**Figure. 2.**
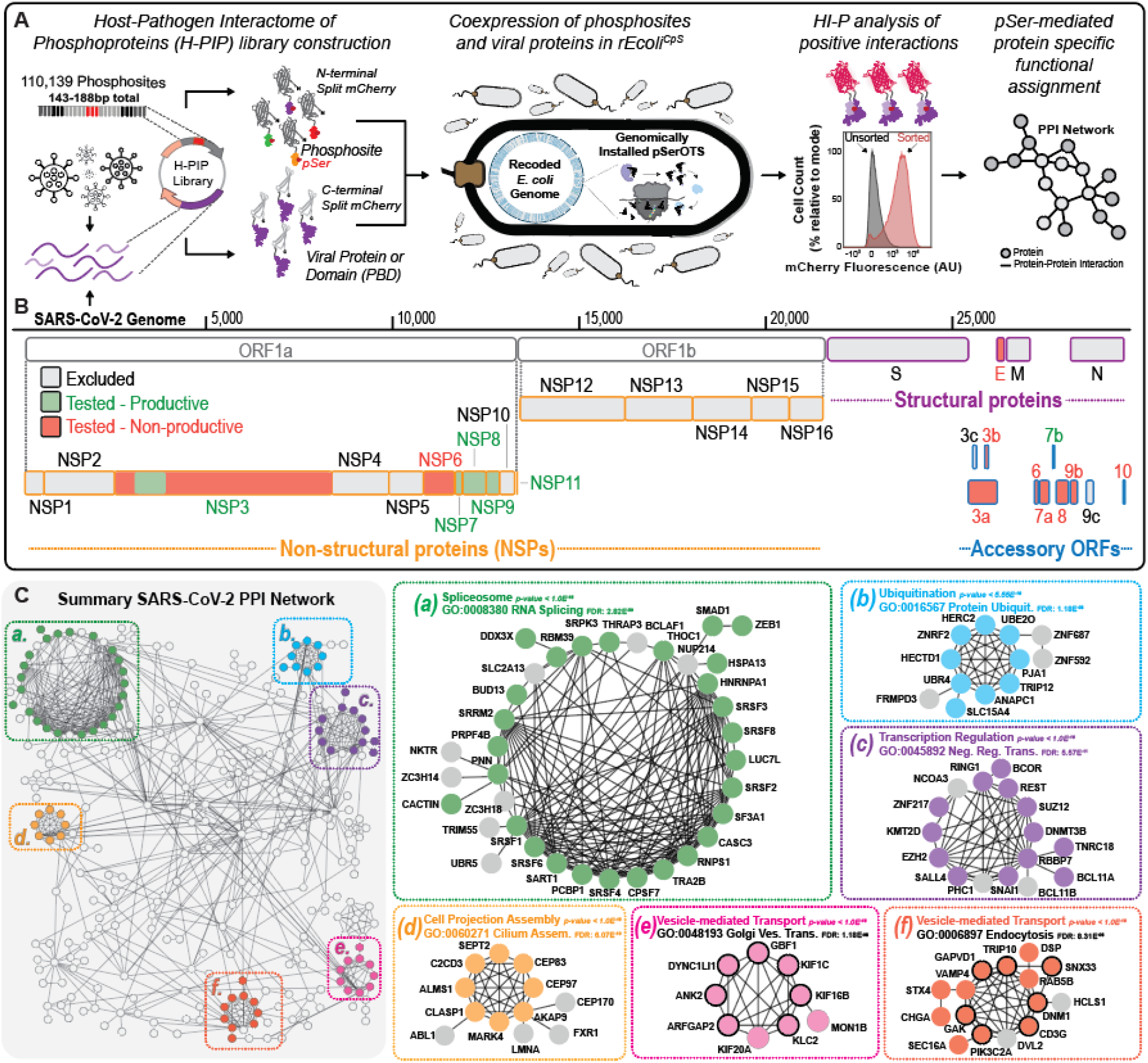
H-PIP workflow and application towards the profiling of SARS-CoV-2 viral protein pSer-mediated PPIs. (**A**) Schematic of H-PIP workflow for the investigation of intracellular pathogen PPIs with host phosphoproteome in genetically modified *E. coli.* (**B**) Representation of SARS-CoV-2 genome highlighting productive (green) or non-productive (red) proteins or domains examined for pSer-specific binding activity. (**C**) Network analysis of high-confidence protein level interactions with greater than 10-fold enrichment. Colors identify subnetworks (a-f): expanded view showing GO-terms and associated false discovery rate (FDR).

Most SARS-CoV-2 structural and major viral replication proteins (e.g. RNA-dependent RNA polymerase) were excluded from analysis and the remaining viral proteins were filtered based on their overall size, known molecular function, and presence of domains of unknown function (DUFs) (Finkel et al., 2021). From our starting pool of fifteen target proteins, our H-PIP analysis yielded system level, phosphosite specific identification of PPIs for six viral proteins. (Figure 2B**; Figure S3. Related to** Figure 2). Collectively, data from the six productive SARS-CoV-2 proteins formed a highly interconnected network made up of functionally enriched subnetworks that overlap well with recent reports of SARS-CoV-2 PPIs (Figure 2C; Table 1**; Data S1**) (Gordon et al., 2020b; Samavarchi-Tehrani et al., 2020; Stukalov et al., 2021). Amongst other host processes, subnetwork interactions were identified for transcription regulation and mRNA processing. Interactions with proteins recruited for vesicle-mediated viral replication and release were also observed, in line with known molecular mechanisms of SARS-CoV-2 (Banerjee et al., 2020). GO enrichment analysis of the biological network highlighted the diverse functional cellular landscape of pSer-dependent PPIs (Raudvere et al., 2019) (**Figure S4; Data S2. Related to** Figure 2). Interactions for biological processes involved in DNA replication, cell cycle regulation, transcription, and cytoskeletal organization were among the most highly enriched. While consistent with results obtained in other studies, our results provide a list of functionally enriched PPIs connected to precise phosphorylation sites across the human proteome and are independent of viral infection stage, cell cycle phase, and cell type.

**Table 1.** Overview of H-PIP PPIs and overlap with external data. Each row in the table represents a viral protein analyzed by H-PIP. The number of pSer-PPIs(*) was determined by identifying pSer-exclusive PPI enrichments greater than 10-fold conserved across replicates. Overlapping SARS-CoV-2 PPIs identified between H-PIP and externally reported networks presented in Gordon et al. AP-MS(†) (2), Stukalov et al. AP-MS(††) (3), and Samavarchi-Tehrani et al. BioID(‡) (9). The presence of domains of unknown function (DUF) and associated molecular functions are also described for each viral protein.

| Protein/ Domain | Num. of pSer-PPIs* | Overlap with AP-MS<sup>†</sup> | Overlap with AP-MS<sup>††</sup> | Overlap with BioID<sup>‡</sup> | Contains DUF? | Associated Function |
| --- | --- | --- | --- | --- | --- | --- |
| NSP3 (AC region) | **90** | No AP-MS | 1 of 2 (Macro) | 16 of 95 | Yes | Involved in viral replication |
| NSP7 | **144** | 11 of 32 | 4 of 6 | 288 of 1043 | No | RNA polymerase complex |
| NSP8 | **186** | 10 of 24 | 3 of 3 | 207 of 1007 | Yes | RNA polymerase complex |
| NSP9 | **145** | 5 of 16 | 9 of 10 | 268 of 1420 | No | Unknown |
| NSP11 | **54** | 0 of 1 | No AP-MS | 7 of 918 | No | Unknown |
| ORF7b | **87** | No AP-MS | 27 of 398 | 35 of 1512 | Yes | Unknown |

ACE2, a primary receptor for viral entry, is expressed in a subset of cells throughout the human body. However, pull-down based PPI studies are often limited to one cell type, which, for studies of SARS-CoV-2, has mostly relied upon cells artificially expressing ACE2 and cells derived from the lung (Leist et al., 2020). H-PIP overcomes this limitation by providing a platform for cell-type independent analysis of viral PPIs. Examination of ACE2 expression profiles revealed many clinically relevant tissues with higher levels of expression when compared to the lung (Figure 3A) (Uhlen et al., 2015). Recent reports from post-mortem examination of COVID-19 patients have described the presence of viral infection in tissues outside the respiratory system, most notably in the heart and testis (Lindner et al., 2020; Ma et al., 2021). Despite these observations, the contribution of SARS-CoV-2 infection of these tissues towards the severity and progression of COVID-19 remains unknown. To provide insight into the role of SARS-CoV-2 infection of tissues outside the lung, we analyzed our set of phosphosite interactions for enrichment of tissue-specific biological processes. One specific process to emerge was spermatogenesis from testicular tissues. Pathology reports from COVID-19 patients have described the impact of viral infection on male reproductive tissues and defects in sperm maturation (Li et al., 2020; Ma et al., 2021). While the underlying mechanisms of this pathology and long-term implications of these findings are unknown, these observations underscore known hormone and sex-dependent discrepancies in patient mortality (Chanana et al., 2020; Samuel et al., 2020). Using protein and phosphosite level PPIs, we identified viral interactions with 48 of the 402 proteins associated with spermatogenesis (Figure 3B). More generally, we observed 126 (of 736) protein interactions across our network currently annotated in the human protein atlas with tissue specific or tissue enhanced expression, illustrating the comprehensive potential of our cell-type independent profiling strategy (Figure 3C).

**Figure 3.**
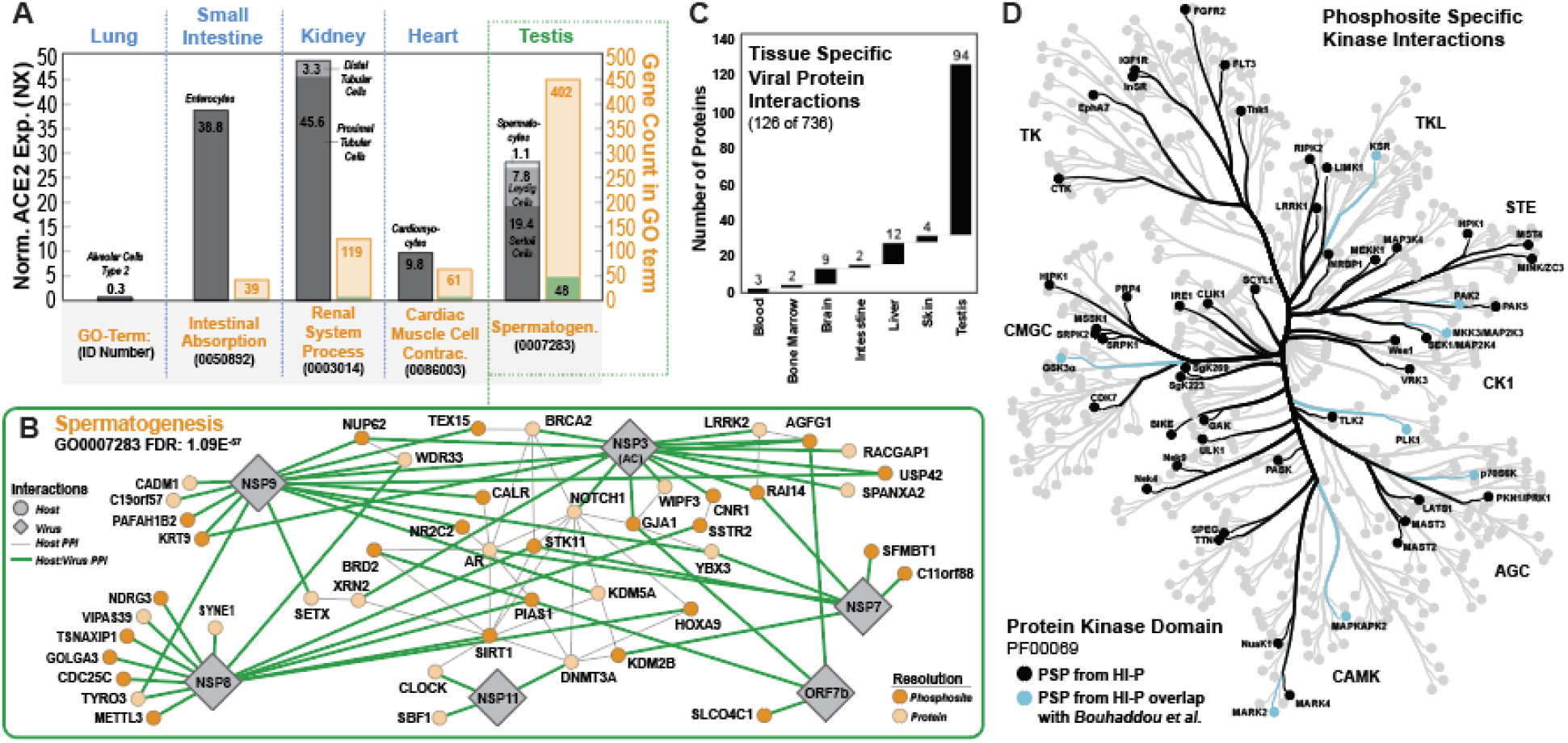
Highly interconnected networks provide access to transient and cell-type independent PPIs. (**A**) Graph of normalized ACE2 expression across specific tissues (left axis, grey bars) in the Human Protein Atlas (HPA) and tissue-type specific biological processes (right axis, orange bars) and overlap with H-PIP (green bar). (**B**) PPI network for proteins involved in spermatogenesis (B); viral protein (diamond) interactions (green lines) and known host PPIs (black lines) at protein (light orange node) and phosphosite-specific (dark orange node) resolution. (**C**) Graph of intersection between viral proteins PPIs from H-PIP and tissue-specific host proteins annotated in HPA. (**D**) Kinome tree depicting phosphosite-specific interactions between viral proteins and PKs (black branches and nodes) and overlap with PKs implicated in previous studies (*4*) (light blue).

Transient PPIs between viral proteins and protein kinases (PKs) are notoriously difficult to capture using traditional methods. As primary effectors of cellular programs, PKs have highly dynamic modification states, cellular localization, and expression levels. H-PIP circumvents these challenges by presenting stably modified PK surfaces for PPI analysis independent of confounding factors in the host cell mentioned above. Our analysis of viral interactions with PKs revealed 52 phosphosite-specific interactions that viral proteins could use to exploit PK function and ultimately lead to cellular reprogramming events to facilitate viral infection and disease progression (Figure 3D). Only eight PKs from our interactome were previously identified in kinase-related studies of SARS-CoV-2 (Bouhaddou et al., 2020). Our list of candidate phosphosite-level PK interactions provides an expanded set of candidate drug targets (many with FDA approved compounds altering target phosphorylation state) that could be explored in future work as a means to mitigate viral infection. As a whole, phosphosite-resolution of interactions across our PPI network provides unprecedented insight into the mechanisms of SARS-CoV-2 infection and pathogenesis in understudied, clinically relevant tissue, ultimately aiding the identification of therapeutic targets and improvement of patient outcomes.

### Secondary viral-human protein complexes suggest discrete but overlapping alternative functions for viral proteins Nsp7 and Nsp8

Beyond their structural roles, limited sets of viral proteins must orchestrate the takeover of cellular machinery to systematically redirect resources towards viral replication and evasion of the host immune response. Accomplishing this necessitates viral proteins often take on multiple roles within an infected cell depending upon the stage of infection (V’Kovski et al., 2021). To varying extent, most SARS-CoV-2 proteins have been observed to interact with host proteins and cellular processes divergent from their ascribed primary functions in viral infection (Wong and Saier, 2021). These expanded interaction profiles implicate alternative functions, particularly for viral protein complexes with pre-defined roles in viral pathogenesis. The primary function for the Nsp7-Nsp8-Nsp12 viral RNA polymerase (RNAP) protein complex, for example, is described as the core complex for RNA-dependent genome replication (Peng et al., 2020). The interactomes of these proteins independent of their complex has been examined through several pull-down based studies, yet a clear picture of their unique roles as individual proteins has yet to emerge. Using H-PIP, we were able to construct PPI networks which recapitulated many of the interactions identified in previous studies (Gordon et al., 2020b). We further expanded the scope and contextual depth of these PPI networks by adding pSer-mediated interactions that implicated functional roles outside of RNA replication (Figure 4A). In addition to pSer-dependent interactions, phosphorylation-independent interactions with the prey library were assessed using a suppressor tRNA (supD) which enables decoding of TAG codons as Ser rather than pSer (**Figure S5**) (Steege, 1983). PPIs identified from the Ser phosphosite interactome allowed us to determine phospho-specificity for interactions observed in the pSer-mediated network and further enhance Nsp7 and Nsp8 network connectivity by including non-phosphorylated phosphosite interactions. The crystal structure for the Nsp7-Nsp8 complex has been reported as a ring structure made up of eight Nsp7-Nsp8 units (Figure 4B) (Peng et al., 2020). Interestingly, the crystal structure for Nsp8 was only able to be solved in the presence of Nsp7 (te Velthuis et al., 2012). In the context of productive PPI networks derived from Nsp8 expression in the absence of Nsp7, this observation suggests an alternative conformation for monomeric Nsp8 that may be intrinsically disordered. This would expose binding surfaces with diverse electrostatic potential, normally sequestered in protein complex, which could contribute to alternative function (Figure 4B).

**Figure 4.**
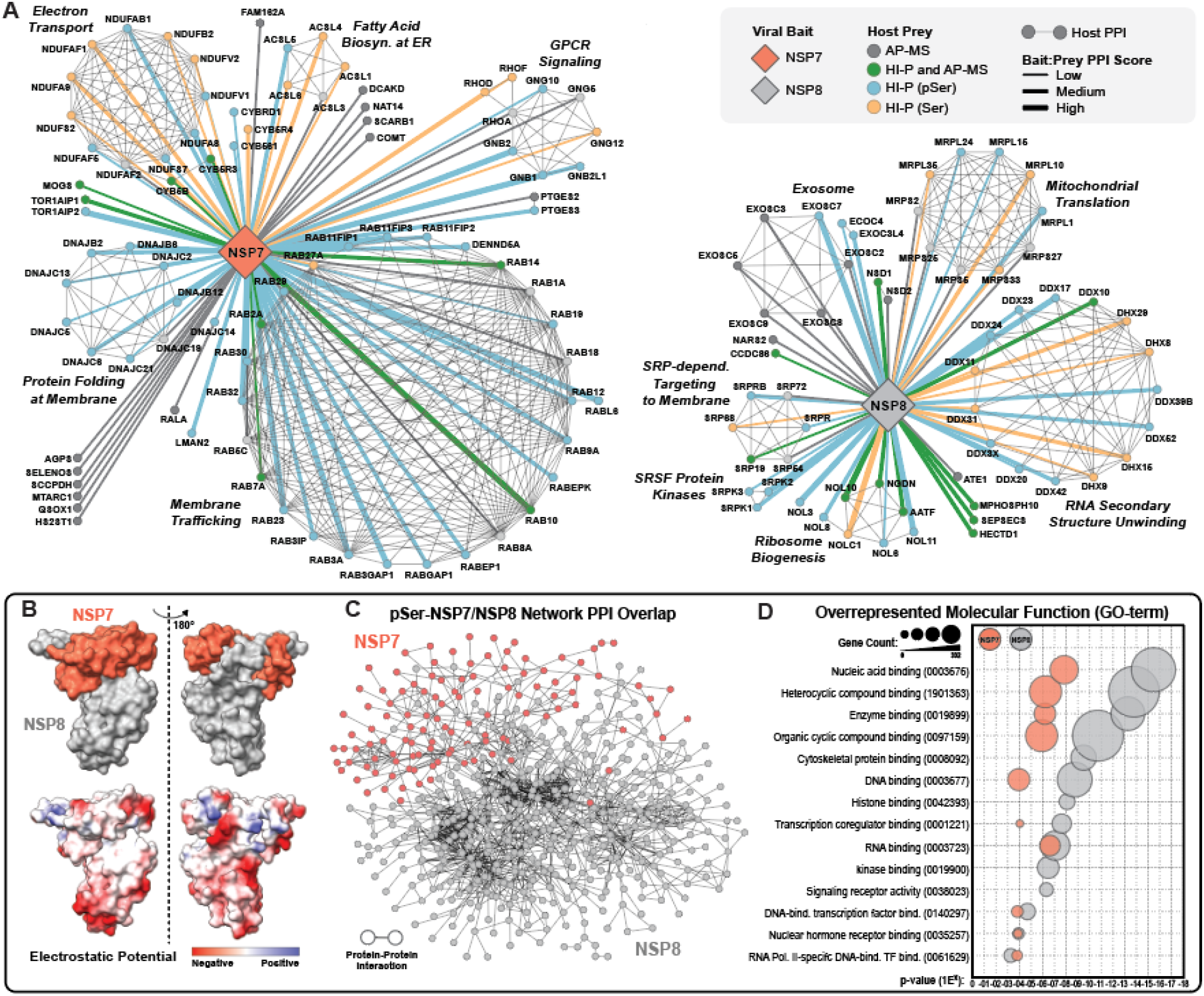
Complex-forming viral proteins have distinct physical interactions but overlapping molecular function. (**A**) Targeted network analysis of Nsp7 and Nsp8 pSer (blue) and Ser (Orange) protein level interactions with low (< 2-fold), medium (2-10 fold), and high (>10-fold) enrichment scores and their intersection (green, overlap) with reported interactions (grey, no overlap). (**B**) Crystal structure of Nsp7-Nsp8 complex displaying surface interactions and electrostatic potential (PDB:7JLT). (**C**) Dynamic network analysis for Nsp7 (red) and Nsp8 (grey) was performed in Cytoscape using DyNet Analyzer. (**D**) Molecular function overrepresentation for proteins in Nsp7 (red) and Nsp8 (grey) networks showing significant enrichments and p-values for overlapping GO-terms.

Dynamic network analysis and superposition of Nsp7 and Nsp8 pSer-mediated PPI networks (expanded from Figure 4A) revealed distinct but highly interconnected protein interaction profiles (Figure 4C). At the functional level, however, GO enrichment analysis of molecular functions in each network revealed a high degree of complementarity both within and across networks (Figure 4D). Consistent with known functional roles, nucleic acid binding was found to be the most significantly enriched molecular function in both networks. Several unique functional enrichments were also observed in each network, most notably for proteins involved in membrane trafficking and kinase binding activity in Nsp7 and Nsp8, respectively. Closer inspection of the Nsp8 PPI network revealed unique pSer-mediated interactions with PKs SRPK1 and SRPK2 (Figure 4A), which are involved in the regulation of mRNA maturation and contribute to mechanisms of virus-mediated cellular reprogramming (Heaton et al., 2020). Taken together, these data implicate a number of alternative functional roles for Nsp7 and Nsp8 in viral pathogenesis independent of their primary structural and enzymatic roles as integral members of the RNAP complex.

### Functionally enriched H-PIP networks support defined host interactions for the viral proteins of unknown function nsp9 and nsp11

Aside from viral proteins possessing well characterized structural or enzymatic properties related to viral propagation, the interactions and functional roles of many viral proteins remains poorly understood. For example, the non-structural proteins nsp9 and nsp11 from SARS-CoV-2 have been shown to interact with host proteins, but the implication of these interactions and contribution to disease progression remains largely unknown (Gordon et al., 2020b).

Nsp9 is a ∼12 kDa NSP with high conservation (98%) to its SARS-CoV homologue. Prior studies of nsp9 from SARS-CoV were targeted at defining its role in as an essential component of viral replication and identified direct single-stranded nucleic acid binding activity and putative interaction with nsp7-nsp8 components of the viral replication and transcription complex proteins (Egloff et al., 2004; Littler et al., 2020). Until recently, little work has been done to identify interactions between nsp9 and the host cellular environment. Pull-down experiments from recombinantly expressed SARS-CoV-2 nsp9 in human cells have revealed a number of interactions with host proteins, yet it its functional role in the alteration of host processes remains ambiguous (Gordon et al., 2020a; Gordon et al., 2020b). To expand the characterization of these interactions and aid in functional assignment, we leveraged H-PIP to identify phospho-specific interactions between nsp9 and the human proteome. Analysis of the sequence-specific H-PIP data revealed a strong binding preference for acidic residues (D/E) at positions −3 to −1 and +1 of the pSer phosphosite (Figure 5A). Interestingly, D/E-rich motif elements have been demonstrated to participate in gene regulation strategies via DNA/RNA mimicry (Banerji et al., 2012). GO analysis of nsp9 binding partners identified significant enrichment for biological processes related to nucleic acid metabolism and genome organization, as well as molecular functions defined by nucleic acid and transcription factor binding (Figure 5B). Assembly of pSer-specific nsp9 interactions generated a highly connected, functionally enriched PPI network centered around host proteins involved in nucleic acid metabolism (Figure 5C). Taken together, the H-PIP data strengthens the role of nsp9 as a key component of viral replication by identifying direct interactions with host nucleic acid processes. These observations support a functional role for nsp9 in the redirection of cellular resources towards viral genome replication via direct phospho-dependent interactions with primary effector proteins of host genome duplication and gene expression.

**Figure 5.**
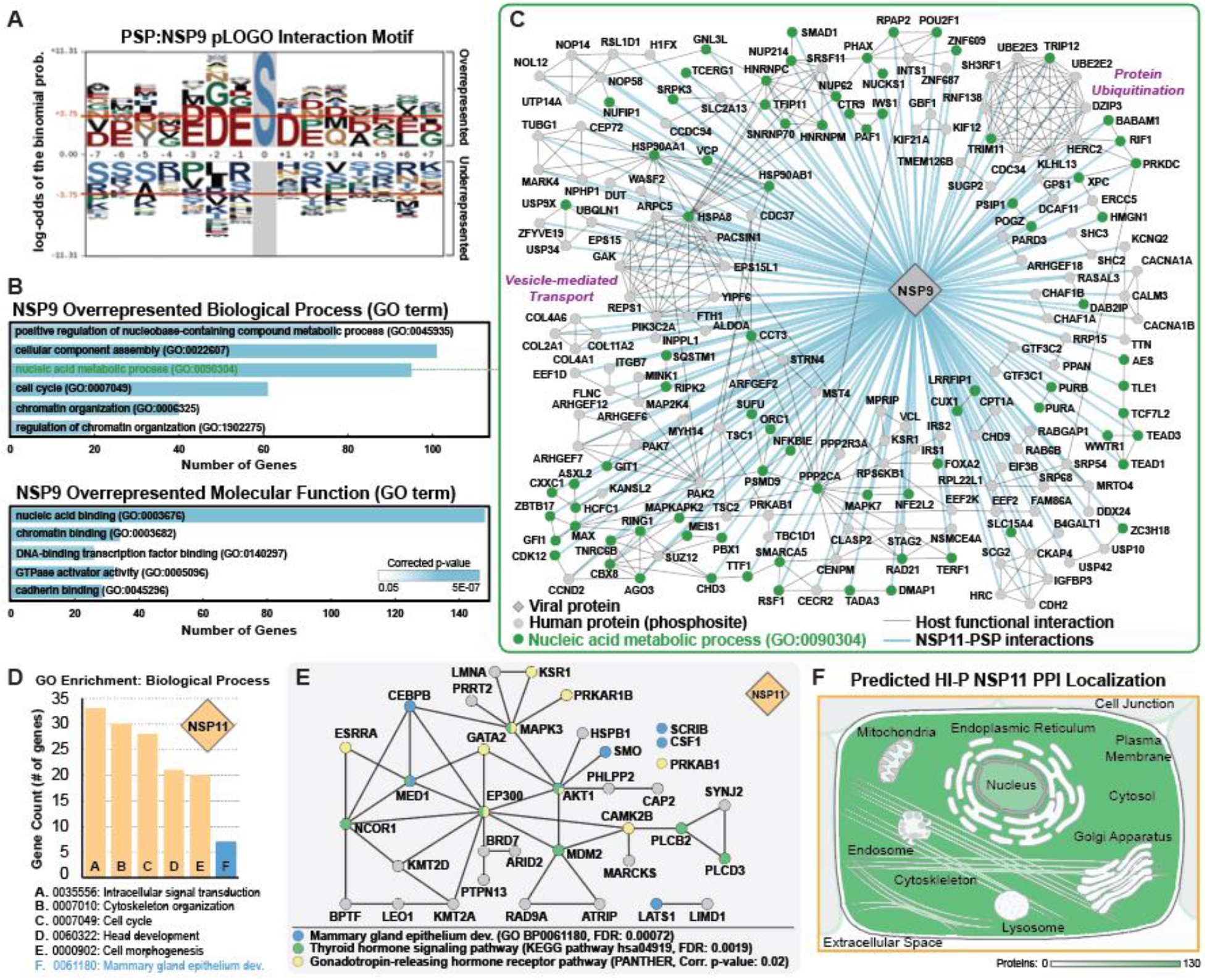
H-PIP indicates putative functional roles in nucleic acid metabolism and hormone regulation for NSP9 and NSP11, respectively. (**A**) pLOGO motif analysis of pSer-specific phosphosites interactions with NSP9. (**B**) PANTHER GO term overrepresentation analysis for biological processes (top) and molecular function (bottom) for pSer-specific NSP9 interactions binned by number of genes in each GO term and corrected p-value. Nucleic acid metabolic process highlighted in green. (**C**) Network analysis of PSP-NSP9 displaying interactions with host PPIs (gray lines) and NSP9 interactions (blue lines) highlighting interactions with genes related to Nucleic acid metabolic process (green nodes). (**D**) PANTHER GO term enrichment analysis for biological processes from pSer-specific interactions with NSP11. (**E**) Subnetwork analysis of NSP11-Host interactions related to mammary gland epithelium dev. (blue nodes), thyroid hormone signaling (green nodes), and Gonadotropin-releasing hormone receptor pathway (yellow nodes). (**F**) Predicted subcellular localization of proteins interacting with NSP11. Color scaled by number of interacting genes in each subcellular compartment.

Mature viral proteins are often produced from proteolytic processing or translational frameshifting events which serve to maximize functional space while minimizing size of the viral genome. Many of these events yield relatively small viral proteins. The 13 amino acid nsp11 protein results from proteolytic cleavage of the pp1a polyprotein at the nsp10/nsp11 junction. Though small in size and intrinsically disordered, nsp11 has been recently observed to possess protein binding capacity (Gadhave et al., 2021; Gordon et al., 2020b). Attempts to characterize nsp11 PPIs within the host have failed to reliably establish a consensus network or functional assignment. Using H-PIP, we assessed the ability of nsp11 to bind PSPs (and their non-phosphorylated equivalents) across the human proteome. Combination of phospho- and non-phsopho-specific PPIs expands the ability of H-PIP to identify putative functional roles across the cellular environment. GO analysis of cumulative protein interactors revealed several biological process enrichments including signal transduction, cellular organization, and mammary gland epithelium development (Figure 5D). Mammary gland tissues upregulate expression of the ACE2 receptor during pregnancy and lactation, making it conditionally susceptible to SARS-CoV-2 infection (Hennighausen and Lee, 2020). Targeted network analysis focused on protein interactors involved in mammary gland development displayed significant pathway enrichments for thyroid signaling and gonadotropin-releasing hormone receptor pathways, both of which have been linked to mammary gland regulation and remodeling (Figure 5E) (Capuco et al., 2008; Ermekova et al., 1997; Russo et al., 1990). Cellular component analysis of host protein interactors was used to determine the hypothetical localization of nsp11 to specific subcellular compartments. In line with biological process and pathway enrichments, localization of nsp11 interactors was predominantly cytoplasmic with significant enrichment at the cytoskeleton and cell-cell junction (Figure 5F). As a whole, the H-PIP derived PPIs implicate a functional, and potentially pathogenic, role for nsp11 in mammary gland tissue through interactions with hormone signaling pathways processes related to cellular morphogenesis. Organism-wide profiling of tissue-specific phospho-dependent viral PPIs is unique to H-PIP and dramatically expands our ability to identify putative functional and pathogenic roles for viral proteins of unknown function.

### ORF7b is a transmembrane SR/RS-binding protein with high SRPK1 phosphosite overlap

Defining systems-level interactions of viral accessory ORFs has proven challenging using traditional interactomic strategies due to their size and often limited biophysical characterization. SARS-CoV-2 accessory ORF7b is an excellent example of the challenges these ORFs present and is often excluded from interactome analyses (Gordon et al., 2020b). Studies that have published interactomes for ORF7b have been difficult to interpret, marked by low study-to-study reproducibility and expansive lists of protein interactions attributed to non-specific binding events. ORF7b is a 5.2 kDa transmembrane protein localized to ER and Golgi membranes during both SARS-CoV and SARS-CoV-2 viral infection (Figure 6A) (Gordon et al., 2020a). To better visualize the features of ORF7b, we constructed biophysical and structural prediction models i*n silico* which yielded a hydrophobic, single transmembrane helix protein model (Figure 6B) with a negatively charged domain localized to the solvent exposed C-terminal end (Figure 6C**, Figure S6**). When compared to the SARS-CoV homologue, the sequence of the transmembrane domain of SARS-CoV-2 ORF7b is highly conserved, while the cytoplasmic protrusion if significantly divergent (Figure 6A).

**Fig. 6.**
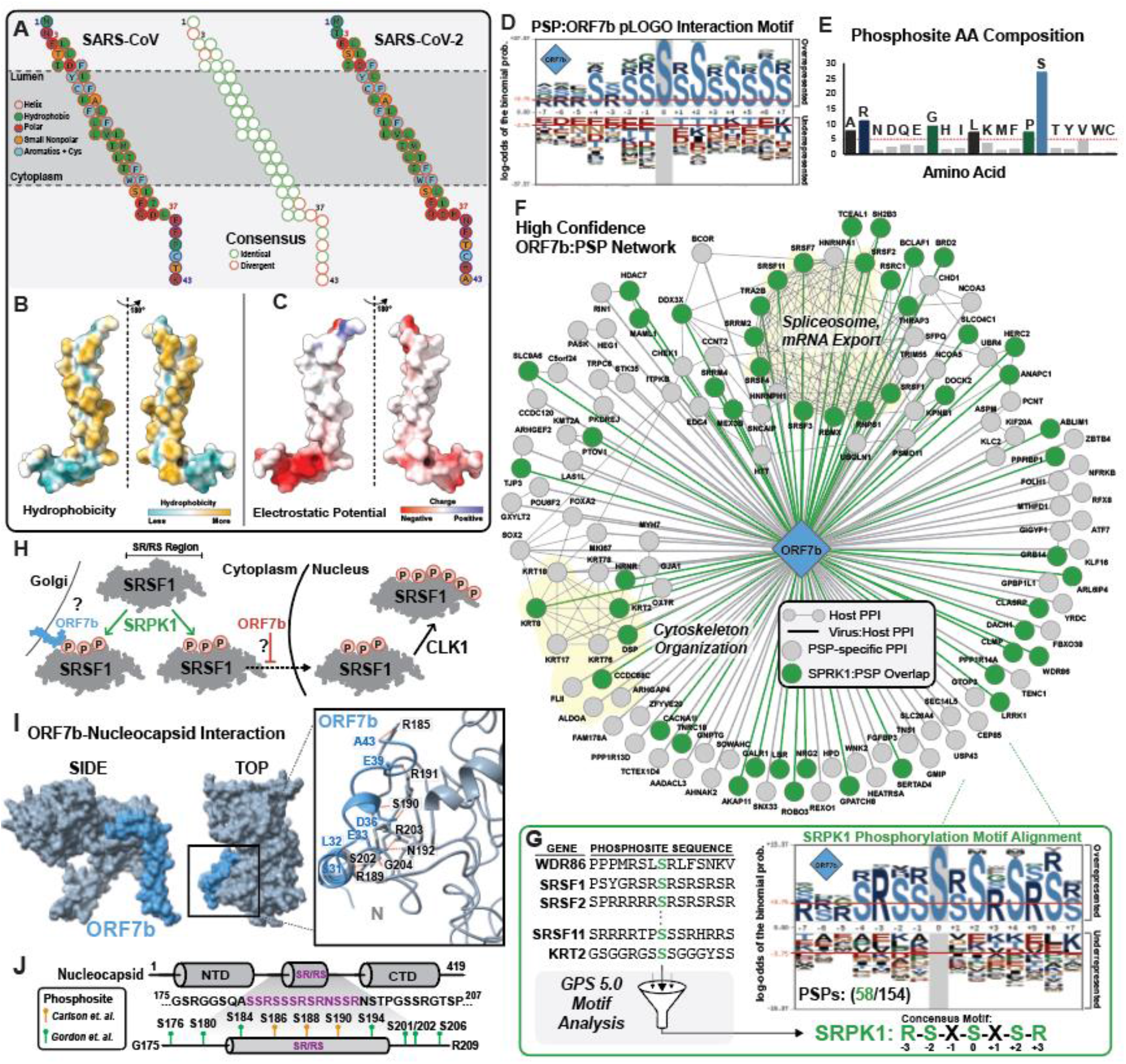
ORF7b is an integral membrane protein with binding preference for SR-rich proteins. (**A**) Transmembrane sequence model for ORF7b displaying helix orientation and amino acid composition color coded by physical property for SARS-CoV (left), SARS-CoV-2 (right), and sequence conservation (middle). (**B**) Structural model for ORF7b determined using Phyre2 structure prediction overlaid with hydrophobicity profile or (**C**) electrostatic potential. (**D**) pLOGO motif analysis of pSer-specific phosphosites interactions with ORF7b. (**E**) Amino acid composition of phosphosites in (D). (**F**) High-resolution PPI network for phosphosite specific interactions with ORF7b and overlap with SRPK1 phosphosites (green). (**G**) SRPK1 phosphorylation site prediction for ORF7b network phosphosites using GPS 5.0. Sequence motif for overlapping phosphosites determined using pLOGO. (**H**) Model for SRSF1 translocation and proposed interference by ORF7b-mediated tethering. (**I**) Structural docking model for ORF7b (blue) with N protein (grey) using HADDOCK. Residue specific protein interactions visualized using Chimera. (**J**) Representation of N protein with SR/RS domain sequence (purple) and compiled previously reported phosphorylation sites (green (*2*), orange (*21*).

We approached the C-terminal region of ORF7b as a domain of unknown function (DUF) and investigated its potential to facilitate pSer-mediated protein binding. Using H-PIP, we determined the ORF7b DUF was able to form productive pSer-mediated protein interactions. Motif enrichment analysis of the productive phosphosite interactions identified a striking binding preference for repetitive S-R rich sequences (Figure 6D-E). Proteins with S-R repeats represent a unique class of proteins whose function and localization are often regulated via phosphorylation of SR/RS domains (Jeong, 2017). These domains are highly enriched in host proteins involved in RNA-related cellular processes, including mRNA maturation and regulation, and have been described as critical to viral replication in diverse viral species (Lai et al., 2009; McFarlane and Graham, 2010; Nikolakaki and Giannakouros, 2020). Using our high resolution phosphosite data, we constructed a pSer-mediated PPI network for ORF7b and identified highly connected subnetworks functionally related to spliceosome-mediated mRNA processing and mRNA export (Figure 6F). Interestingly, one of the primary mediators of SR/RS phospho-regulation is SRPK1, a PK strongly implicated as an essential mediator of viral pathogenesis (Heaton et al., 2020; Tunnicliffe et al., 2019). We interrogated the ORF7b phosphosite network for SRPK1 phosphorylation sites using GPS 5.0 and found a high degree of overlap (58 of 154) between phosphosites identified by H-PIP and predicted SRPK1 phosphorylation sites (Figure 6F-G). Motif analysis of the SRPK1-mediated phosphosite sub-pool was consistent with known consensus recognition motifs of the kinase (Figure 6G). The significant overlap observed between ORF7b PPIs and SRPK1 phosphorylation sites implicates ORF7b as a central viral mediator of perturbations to SRPK1-directed host RNA metabolism. As an integral membrane protein, one potential mechanism of ORF7b-mediated cellular reprogramming may be its ability to alter host protein localization by physically tethering proteins to a host membrane. Serine and Arginine Rich Splicing Factor 1 (SRSF1), for example, is a nuclear splicing factor functionally regulated through protein localization mediated by SRPK1-depenedent phosphorylation of its SR/RS domain. Phosphorylation of SRSF1 triggers nuclear import from the cytoplasm and subsequent phosphorylation by PK CLK1 (Aubol et al., 2013). We observed multiple phosphosite-specific interactions between ORF7b and phosphorylated SR/RS-domain of SRSF1 which could enable tethering of SRSF1, thus preventing nuclear translocation and contributing to global disruption of mRNA splicing observed in infected cells (Figure 6H) (Banerjee et al., 2020). At the same time, host proteins involved in RNA metabolism, including SRSF1, have been shown to physically associate with viral RNA (Flynn et al., 2021). Tethering via golgi membrane-bound ORF7b interactions would re-localize these host factors to participate in viral RNA processing events within the neighboring viral replication transcription complex (RTC), potentially making ORF7b essential to recruitment of host factors during viral genome replication.

In addition to host proteins, the SARS-CoV-2 nucleocapsid (N) protein contains an SR/RS domain localized to the disordered region between the N- and C-terminal domains that has been demonstrated to be a critical determinant of efficient viral replication (Figure 6I-J) (Tylor et al., 2009). The primary function of the N protein is packaging the viral genome during virion assembly. Association of N with viral RNA is phosphorylation-state dependent and regulated through recruitment of host phosphatases and PKs, including SRPK1 (Carlson et al., 2020). In the early infection phase, N:viral RNA complex dissociation is primed by SRPK1-mediated phosphorylation of the SR/RS domain. In later phases of infection, N protein is dephosphorylated in preparation for viral RNA packaging during virion assembly, however the exact mechanism by which phosphorylated N protein is recruited from its alternative cytoplasmic functions and re-localized to the RTC is unknown. One potential mechanism of N protein recruitment is through interactions between ORF7b and phosphosites within the N SR/RS region. Since a complete structure of SARS-CoV-2 N protein has yet to be reported, we constructed a theoretical model using *in silico* structure prediction. Using this structure, we compiled structural docking models with ORF7b and found that the SR/RS region of N protein is accessible to, and able to form interactions with, the exposed cytoplasmic domain of ORF7b (Figure 6I**; Figure S7-S8. Related to** Figure 6). This mechanism is further supported by reported co-localization of ORF7b and the RTC to the ER/golgi membrane and by observations that the ORF7b homolog from SARS-CoV is one of the few viral proteins packaged within a mature viral particle (Gordon et al., 2020a; Schaecher et al., 2007).

## Discussion

Our approach constitutes a high-resolution strategy to identify physical interactions across the human phosphoproteome that could be exploited by intracellular pathogens to reprogram the host cell during infection. H-PIP bypasses many of the limitations of traditional protein interaction analyses by removing PPI determination from the host cellular environment where interactions may be cell-type dependent or regulated temporally and spatially by infection-mediated cellular alterations. More importantly, H-PIP analysis uniquely provides important data regarding pSer-mediated PPI specificity and precise pSer site identity missing from current PPI data. Another unique feature of H-PIP is its ability to capture transient PPIs due to the intrinsic irreversibility of fluorophore maturation initiated upon protein contact.

Cumulative analysis of viral PPIs at phosphosite resolution provides a unique opportunity to identify phosphorylation-mediated interactions and pair them with known PK recognition motifs. Together with contextual information obtained from functional enrichments within viral PPI networks, these data could be used to direct novel therapeutic strategies aimed at counteracting PK-mediated host-pathogen reprogramming. Subsets of interaction data can be combined for proteins in complex and used to expand alternative functional roles. For individual viral proteins, our approach generates a unique cross-section of viral mechanisms mediated by protein phosphorylation that can be used to assign putative functional roles to proteins or domains of unknown function. As a whole, H-PIP is a complementary approach to traditional PPI determination that provides high-resolution phosphorylation-centric context to underlying mechanisms of pathogenesis and disease progression for intracellular pathogens.

## Acknowledgments

We would like to acknowledge members of the Rinehart and Isaacs lab for insightful discussion during the preparation of this manuscript.

## Author contributions

Conceptualization: KM, JR

Methodology: KM, SR, JR

Investigation: KM, JM, SR, JR

Visualization: KM, JM

Funding acquisition: KM, JR

Project administration: KM, JR

Writing – original draft: KM, JR

Writing – review & editing: KM, JM, JR

## Data and materials availability

The mass spectrometry proteomics data have been deposited to the ProteomeXchange Consortium (http://proteomecentral.proteomexchange.org) via the PRIDE partner repository with the data set identifier PXD027537. High-throughput sequencing raw data files have been uploaded to the Sequence Read Archive (BioProject ID PRJNA749552).

## Material and Methods

### General Molecular Biology

All DNA oligonucleotides with standard purification and desalting and Sanger DNA sequencing services were obtained through the Keck DNA Sequencing facility and Oligonucleotide Resource, Yale University. Unless otherwise stated, all cultures were grown at 37 °C in LB-Lennox medium (LB, 10 g/L bacto tryptone, 5 g/L sodium chloride, 5 g/L yeast extract). LB agar plates were LB plus 20 g/L bacto agar. Antibiotics were supplemented for selection, where appropriate (Kanamycin, 50 μg/mL and Ampicillin, 100 μg/mL). *E. coli* Top10 cells (Invitrogen, Carlsbad, CA) were used for cloning and plasmid propagation. NEBuilder HiFi Assembly Mix and restriction enzymes were obtained from New England BioLabs. Plasmid DNA preparation was carried out with the QIAprep Spin Miniprep Kit (Qiagen).

A complete list of strains and plasmids may be found in **Supplemental Table 2**. All plasmids were transformed into recipient strains by electroporation. Electrocompetent cells were prepared by inoculating 20 mL of LB with 200 μL of saturated culture and growing at 37 °C until reaching an OD_600_ of 0.4. Cells were harvested by centrifugation at 8,000 RPM for 2 min. at 4 °C. Cells pellets were washed twice with 20 mL ice cold 10% glycerol in deionized water (dH_2_O). Electrocompetent pellets were re-suspended in 100 μL of 10% glycerol in dH2O. 50 ng of plasmid was mixed with 50 μL of re-suspended electrocompetent cells and transferred to 0.1 cm cuvettes, electroporated (BioRad GenePulser™, 1.78 kV, 200 Ω, 25 μF), and then immediately resuspended in 600 μL of LB. Transformed cells were recovered at 37 °C for one hour and 100 μL was subsequently plated on appropriate selective medium.

### Plasmid Construction

To create OTS component integration plasmids, pSerRS (SepRS9) from SepOTSλ (Addgene # 68292) and a recoded EF-pSer21 synthesized as a gBlock (IDT) were amplified and placed under the control of the glnS*, TRC*, proD, or OXB20 promoter (where indicated) and assembled between the FRT sites of plasmid pKD4 by gibson assembly using NEBuilder HiFi DNA Assembly master mix (NEB). 2x tRNA cassettes integration plasmids were constructed from a 2x-tRNA construct under the control of a proK tRNA promoter and terminator and separated by a valU tRNA linker that was ordered as a DNA fragment from Genewiz. tRNA cassettes were likewise assembled into a pNAS1B backbone by gibson assembly. H-PIP plasmids (prior to library introduction) were constructed by amplifying E. coli codon optimized SARS-CoV-2 genes from plasmid vectors synthesized by GinkoBioworks and distributed by Addgene. Amplified genes were digested with NdeI and SacI, ligated into the background H-PIP plasmid, and transformed into Top10 chemically competent cells to enable selection of ligated clones on LB+Kanamycin. Following selection and verification by DNA sequencing, the pSer phosphosite library was cloned into each H-PIP plasmid, as described previously (Barber et al., 2018).

### rEcoli^CpS^ Strain Construction

Strain modification using a lambda red based strategy was performed as previously described (Datsenko and Wanner, 2000). Briefly, transformants carrying a lambda red plasmid (pKD46) were grown in 20 mL LB cultures with ampicillin and l-arabinose at 30°C to an OD_600_ of ≈0.6 and then made electrocompetent. 10–100 ng of PCR product amplified from integration plasmids was transformed into the prepared cells and recovered in 1 mL of LB for 1 h at 37°C. Following recovery, one-half was spread onto LB-agar plates with Kanamycin for selection. Gene deletion was verified by PCR and selected colonies were purified non-selectively at 37°C remove pKD46. The KAN deletion cassette was excised from the integration locus using FLP recombinase (pCP20).

### H-PIP Analysis

*Fluorescence-activated cell sorting for H-PIP:* 20 mL of rEcoli^CpS^ cells were grown to OD_600_ of 0.4 and prepared for electrotransformation as described above. Prepared cells were transformed with 1 μL of SARS-CoV-2 H-PIP library plasmid (approximately 100 ng/μL). The cells were then resuspended in 1.2 mL of S.O.C. medium (Thermo) and incubated for 1 h at 37 °C and 230 rpm in a 15 mL culture tube. Following outgrowth, cells were then transferred to 20 mL of LB with 100 ng/μL ampicillin in a 50 mL conical tube and grown overnight at 37 °C and 230 rpm.

The next morning, cultures were diluted to an OD_600_ of 0.15 in 5 mL of LB containing 100 ng/μL ampicillin and grown at 37 °C and 230 rpm. The cells were grown until OD_600_ reached mid-log (0.6-0.8), then protein expression was induced using 0.2 % arabinose and 100 ng/μL anhydrotetracycline, and grown at either 37 °C for 3 hours followed by 20 °C for hours, or 20 °C for 20 hours only. Following induction, 100 μL of cells were diluted in 3 mL ice cold PBS in a 5 mL polystyrene tube (Falcon).

Using a BD FACSAria III, cells were interrogated for mCherry-based fluorescence using a 561-nm laser. 50,000 cells were sorted using a gate empirically determined to yield substantially enriched fluorescent signal in regrown cell populations, which differed for each phospho-binding domain (generally 100-200 cells/s from right tail). Cells were sorted directly into 2 mL LB without antibiotic, recovered at 37 °C and 230 rpm for 2 h, and then transferred to a 50 mL conical tube containing 20 mL LB with 100 ng/μL ampicillin. After growth overnight, the procedure for protein expression, preparation for FACS, and cell sorting was repeated, using the same sorting and gating parameters as the first round of sorting. Cells were then recovered, regrown, induced and prepared for FACS as above. Cellular mCherry fluorescence was then observed using the FACSAria III. Plasmid libraries isolated by miniprep of twice-sorted cell populations were prepared for next-generation sequencing as described below.

#### High-throughput sequencing information

PCR amplicons of phosphosite gene libraries in the modified pNAS1B vector for H-PIP experiments were generated by PCR. DNA amplicons were isolated using a BluePippin with size selection range of 100-200bp. Following size selection, amplicons were purified using a PCR clean-up kit (Qiagen) and submitted to Mass. General Hospital (MGH) DNA sequencing facility for 250pb illumina paired-end read sequencing.

Sequencing reads were first filtered for quality using Trimmomatic33, which applied a sliding window filter of width 2 bp and a Phred score cutoff of 30. If the average quality score over two consecutive bases fell below 30, the read was trimmed to remove the remaining bases. Quality trimmed read pairs were then merged using BBMerge with the stringency set to “strict” (sourceforge.net/projects/bbmap). Using custom scripts, the merged reads were then sorted and assigned to the various input libraries based on barcodes added during the PCR amplification step. The variable sequence region for each amplicon was then extracted and for each input library the abundance of every unique sequence was calculated. In order to determine library coverage, sequencing reads were filtered for quality using Trimmomatic with a sliding window filter of 2bp and Phred score cutoff of 30. Additionally, the first 5bp were trimmed from the start of the reads. Subsequently, the trimmed read pairs were merged using BBMerge with the stringency set to “strict”. The FASTQ file of merged read pairs was then aligned to a FASTA file containing each of the library member sequences using the BWA-mem algorithm with the –M option. The resultant alignment files were then sorted and indexed using samtools and the mappings to each library member were evaluated using BBMap’s pileup.sh with “secondary=false”.

### Phostag SDS-PAGE and Immunoblotting

100 μM Phos-tag™ acrylamide (Wako) within hand-cast 12% acrylamide gels were used for separation of phosphorylated reporter proteins. SDS-PAGE gels (4–15% acrylamide, Bio-Rad) and Phos-tag™ gels were transferred onto PVDF membranes. All gels were visualized by immunoblot. Anti-His immunoblots were performed using 1:2,500 diluted rabbit Anti-6xHis antibody (PA1-983B, Thermo Fisher Scientific) in 5% w/v milk in TBST for 1 h. Secondary antibody incubations used 1:10,000 diluted donkey anti-rabbit HRP (711-035-152, Jackson ImmunoResearch) in 5% w/v milk in TBST for 1 h. Protein bands were then visualized using Clarity ECL substrate (Bio-Rad) and an Amersham Imager 600 (GE Healthcare Life Sciences).

### Protein Purification

#### MS-READ reporter purification

Frozen *E. coli* cell pellets were thawed on ice and pellets were lysed by sonication with lysis buffer consisting of 50 mM Tris-HCl (pH 7.4, 23°C), 500 mM NaCl, 0.5 mM EGTA, 1mM DTT, 10 % glycerol, 50 mM NaF, and 1 mM Na_3_O_4_V. The extract was clarified with two rounds of centrifugation performed for 20 minutes at 4 °C and 14,000 x g. Cell free extracts were applied to Ni-NTA metal affinity resin and purified according to the manufacturer’s instructions. Wash buffers contained 50 mM Tris pH 7.5, 500 mM NaCl, 0.5 mM EGTA, 1mM DTT, 50 mM NaF, 1 mM Na_3_VO_4_ and increasing concentrations of imidazole 20 mM, 40mM, and 60mM, sequentially. Proteins were eluted with wash buffer containing 250 mM imidazole. Eluted protein was subjected to 4 rounds of buffer exchange (20mM Tris pH 8.0 and 100mM NaCl) and concentrated using a 30 kDa molecular weight cutoff spin filter (Amicon).

### Mass Spectrometry

#### MS-READ analysis

Affinity purified, buffer exchanged protein was digested and analyzed by mass spectrometry as described previously with some modifications. Briefly, the concentration of protein was determined by UV280 spectroscopy and 5 μg ELP-GFP (MS-READ) reporter from *E. coli* was and dissolved in 12.5 μl solubilization buffer consisting of 10 mM Tris-HCl pH=8.5 (23°C), 10 mM DTT, 1 mM EDTA and 0.5 % acid labile surfactant (ALS-101, Protea). Samples were heat denatured for 20 min at 55 °C in a heat block. Alkylation of cysteines was performed with iodoacetamide (IAM) using a final IAM concentration of 24 mM. The alkylation reaction proceeded for 30 min at room temperature in the dark. Excess IAM was quenched with DTT and the buffer concentration was adjusted using a 1 M Tris-HCl pH 8.5 resulting in a final Tris-HCl concentration of 150 mM. The reaction was then diluted with water and 1 M CaCl_2_ solution to obtain a ALS-101 concentration of 0.045 % and 2 mM CaCl_2_ respectively. Finally, sequencing grade porcine trypsin (Promega) was added to obtain an enzyme/protein ratio of 1/5.3 and the digest was incubated for 15 h at 37 °C without shaking. The digest was quenched with 20% TFA solution resulting in a sample pH of 2. Cleavage of the acid cleavable detergent proceeded for 15 min at room temperature. Digests were frozen at −80 °C until further processing. Peptides were desalted on C_18_ UltraMicroSpin columns (The Nest Group Inc.) essentially following the instructions provided by the manufacturer but using 300 μl elution solvent consisting of 80% ACN, 0.1% TFA for peptide elution. Peptides were dried in a vacuum centrifuge at room temperature. Dried peptides were reconstituted and analyzed by LC-MS/MS.

#### Digestion of intact E. coli for shotgun proteomics

20 mL cultures were inoculated to a starting OD 600nm of 0.01 in LB media using an overnight culture to stationary phase. After reaching mid-log, cells chilled on ice and pelleted by centrifugation for 2 min at 8000 rpm. The resulting pellet was frozen at −80 °C for downstream processing. For cell lysis and protein digest, cell pellets were thawed on ice and 2 ul of cell pellet was transferred to a microcentrifuge tube containing 40 μl of lysis buffer (10 mM Tris-HCl pH 8.6, 10 mM DTT, 1 mM EDTA, and 0.5 % ALS). Cells were lysed by vortex for 30 s and disulfide bonds were reduced by incubating the reaction for 30 min. at 55 °C. The reaction was briefly quenched on ice and 16 μl of a 60 mM IAM solution was added. Alkylation of cysteines proceeded for 30 min in the dark. Excess IAM was quenched with 14 μl of a 25 mM DTT solution and the sample was then diluted with 330 μl of 183 mM Tris-HCl buffer pH 8.0 supplemented with 2 mM CaCl_2_. Proteins were digested overnight using 12 μg sequencing grade trypsin. Following digestion, the reaction was then quenched with 12.5 μl of a 20 % TFA solution, resulting in a sample pH<3. Remaining ALS reagent was cleaved for 15 min at room temperature. The sample (∼30 μg protein) was desalted by reverse phase clean-up using C_18_ UltraMicroSpin columns. The desalted peptides were dried at room temperature in a rotary vacuum centrifuge and reconstituted in 30 μl 70 % formic acid 0.1 % TFA (3:8 v/v) for peptide quantitation by UV_280_. The sample was diluted to a final concentration of 0.2 μg/μl and 5 μl (1 μg) was injected for LC-MS/MS analysis.

#### Data acquisition and analysis

LC-MS/MS was performed using an ACQUITY UPLC M-Class (Waters) and Q Exactive Plus mass spectrometer. The analytical column employed was a 65-cm-long, 75-μm-internal-diameter PicoFrit column (New Objective) packed in-house to a length of 50 cm with 1.9 μm ReproSil-Pur 120 Å C18-AQ (Dr. Maisch) using methanol as the packing solvent. Peptide separation was achieved using mixtures of 0.1% formic acid in water (solvent A) and 0.1% formic acid in acetonitrile (solvent B) with either a 90-min gradient 0/1, 2/7, 60/24, 65/48, 70/80, 75/80, 80/1, 90/1; (min/%B, linear ramping between steps). Gradient was performed with a flowrate of 250 nl/min. At least one blank injection (5 μl 2% B) was performed between samples to eliminate peptide carryover on the analytical column. 100 fmol of trypsin-digested BSA or 100 ng trypsin-digested wildtype K-12 MG1655 *E. coli* proteins were run periodically between samples as quality control standards. The mass spectrometer was operated with the following parameters: (MS1) 70,000 resolution, 3e6 AGC target, 300–1,700 m/z scan range; (data dependent-MS2) 17,500 resolution, 1e^6^ AGC target, top 10 mode, 1.6 m/z isolation window, 27 normalized collision energy, 90 s dynamic exclusion, unassigned and +1 charge exclusion. Data was searched using Maxquant version 1.6.10.43 (Cox and Mann, 2008) with Deamidation (NQ), Oxidation (M), and Phospho(STY) as variable modifications and Carbamidomethyl (C) as a fixed modification with up to 3 missed cleavages, 5 AA minimum length, and 1% FDR against a modified Uniprot *E. coli* database containing custom MS-READ reporter proteins. MS-READ search results were analyzed using Skyline version 20.1.0.31 and proteome search results were analyzed with Perseus version 1.6.2.2.

### Bioinformatic Analyses

#### Phosphosite Motif Analysis

Phosphosite motif analysis was conducted using pLOGO v1.2.0 probability logo generator (www.plogo.uconn.edu) (O’Shea et al., 2013). Each phosphosite sequence from the enriched population was trimmed to central pSer +/-7 amino acids and uploaded to pLOGO for alignment. For statistical enrichment analysis, a background library of all potential phosphosites was included. Significantly enriched positions have p < 0.05.

#### GO Enrichment Analysis

GO enrichment analysis was performed using both g:Profiler and PANTHER analysis software. An input list of proteins (from aggregated enriched PPI) was submitted to the tool g:GOSt of g:Profiler (version e103_eg50_p15_68c0e33) with g:SCS multiple testing correction method applying significance threshold of 0.05 (Raudvere et al., 2019). Gene set analysis, functional classification, and statistical overrepresentation testing for individual viral proteins was conducted using PANTHER v16.0 (www.Pantherdb.org). Statistical overrepresentation testing for GO biological process and molecular function was assessed using Fisher’s Exact test using all genes in human reference gene set to calculate FDR (Mi et al., 2013).

#### Network Construction and Visualization

Preliminary network construction to assign known host PPIs was facilitated using StringDB v11 (www.string-db.org) (Szklarczyk et al., 2019). Input proteins from enriched networks were analyzed for connectivity using only experimental and database derived interaction sources (BIND, DIP, GRID, HPRD, IntAct, MINT, and PID databases). Network connectivity was exported to Cytoscape for further analysis. Network visualization and construction was performed in Cytoscape v3.8.2 (www.cytoscape.org) (Shannon et al., 2003).

#### Structure Prediction and Molecular Docking

Structure prediction for SARS-CoV-2 ORF7b and full-length Nucleocapsid protein was conducted using the Phyre2 v2.0 (http://www.sbg.bio.ic.ac.uk/phyre2) protein fold recognition software in intensive modeling mode as described in (Kelley et al., 2015). The resulting structures were downloaded and used to enable molecular docking simulations using the HADDOCK v2.4-2021.05 server (www.wenmr.science.uu.nl/haddock2.4/) (van Zundert et al., 2016). The HADDOCK simulation was run using default parameters except for specification of predicted protein interfaces with ambiguous interaction restraints between the C-terminal domain of ORF7b and SR/RS domain of the Nucleocapsid. Top clusters were assessed based on HADDOCK score, van der Waal energy, and electrostatic energy calculated from iterative rounds of simulation and refinement. Docking structure were visualized using UCSF ChimeraX (Goddard et al., 2018).

#### Kinase Substrate Prediction

Putative substrates for SRPK1 were identified using Group-based Prediction System (GPS) v5.0 (Wang et al., 2020). Amino acid sequences corresponding to phosphosites from enriched H-PIP networks were provided as input and kinase motifs were filtered (Serine-Threonine Kinase/CMGC/SRPK/SRPK1). “High” threshold phosphorylation sites were assessed for phosphosite overlap with H-PIP.

