## Supplemental Data and Figures for "Cell type-independent profiling of interactions between intracellular pathogens and the human phosphoproteome"

**This PDF file includes:**

Figs. S1 to S8
Tables S1
Captions for Data S1 to S2

**Other Supplementary Materials for this manuscript include the following:**

Data S1 to S2 [S1: HI-P NGS Supplementary Data Summary]
[S2: GO Enrichment Analysis Supplementary Data Summary]

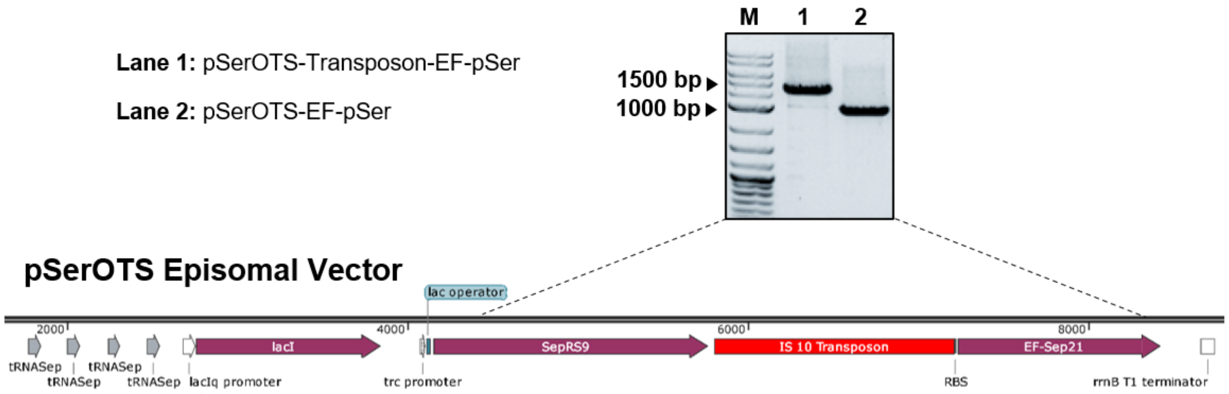


**Figure S1. Transposon insertion in episomal pSerOTS.** PCR amplification of the pSerRS-EF-pSer operon indicated an insertion which was confirmed by sanger sequencing as an IS10 transposon integration event located at the end of pSerRS (SepRS9).

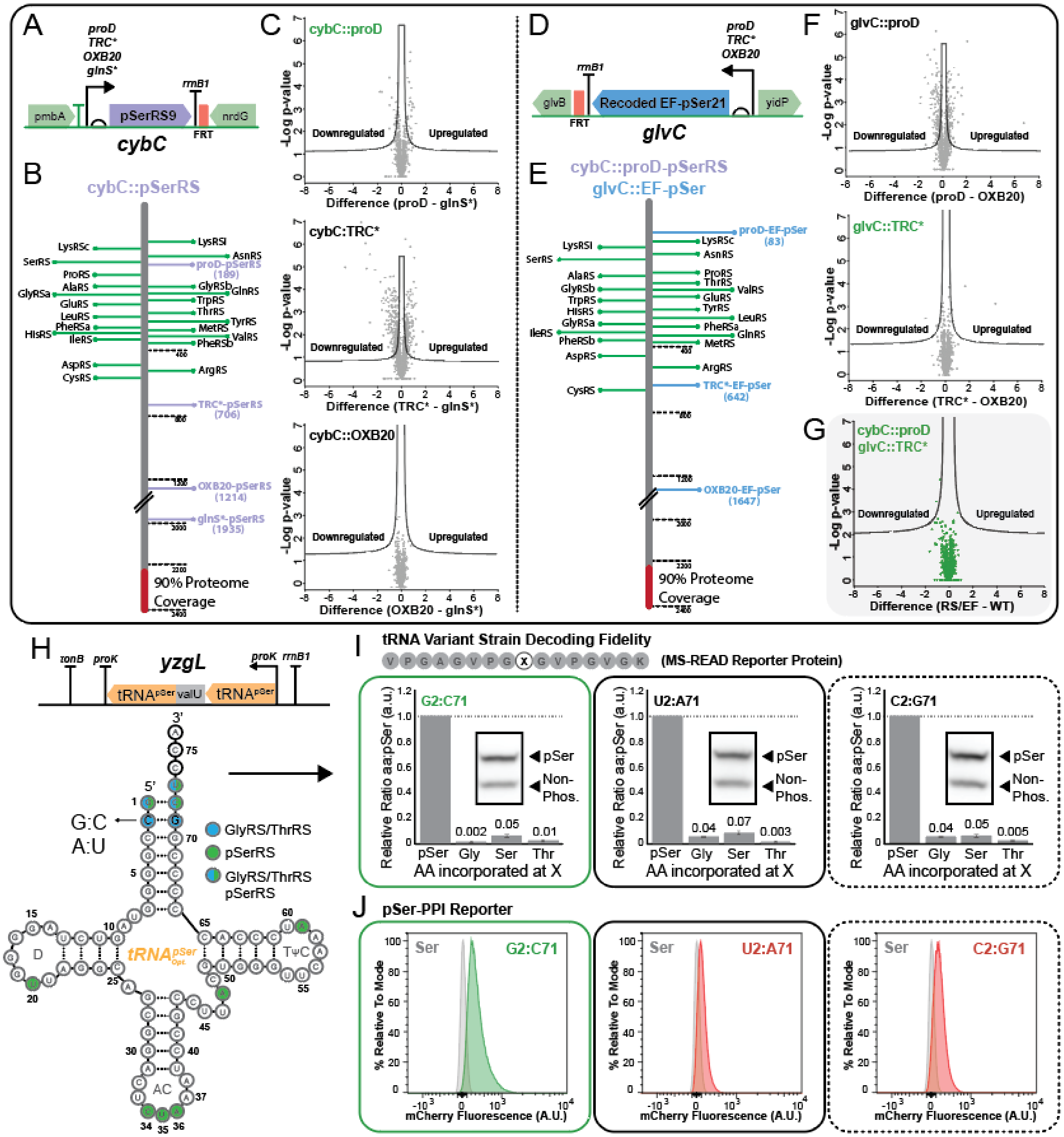
A
B
C
D
E
F
G
H
I
J
cybC
glvC
cybC::pSerRS
cybC::proD-pSerRS
glvC::EF-pSer
cybC::proD
cybC::TRC*
cybC::OXB20
glvC::proD
glvC::TRC*
90% Proteome Coverage
Downregulated
Upregulated
-Log p-value
Difference (proD - glnS*)
Difference (TRC* - glnS*)
Difference (OXB20 - glnS*)
Difference (proD - OXB20)
Difference (TRC* - OXB20)
Difference (RS/EF - WT)
Recoded EF-pSer21
yzgL
GlyRS/ThrRS
pSerRS
GlyRS/ThrRS pSerRS
tRNA Variant Strain Decoding Fidelity
(MS-READ Reporter Protein)
G2:C71
U2:A71
C2:G71
Relative Ratio aa:pSer (a.u.)
AA incorporated at X
pSer-PPI Reporter
% Relative To Mode
mCherry Fluorescence (A.U.)

**Figure S2. Integration and characterization of pSerOTS components.** (**A**) Schematic of pSerRS promoter variants at cybC locus. (**B**) Quantification of proteins in the *E. coli* proteome by mass spectrometry from cells with integrated pSerRS variants, relative to expression levels for native aminoacyl-tRNA synthetases (Green). Relative protein expression was calculated by averaging label-free quantification scores (iBAQ) obtained from MaxQuant analysis(*33*), n=3. (**C**) Volcano plot depicting changes in protein expression between integrated pSerRS variant (OXB20, TRC*, proD) strains and cells with glnS*-pSerRS. Proteins above the curved lines represent significant expression changes; n = 3. (**D**) Schematic of EF-pSer promoter variants at glvC locus. (**E**) Quantification of proteins in the *E. coli* proteome by mass spectrometry from cells with integrated EF-pSer variants, relative to expression levels for native aminoacyl-tRNA synthetases (Green). Relative protein expression was calculated by averaging label-free quantification scores (iBAQ) obtained from MaxQuant analysis(*33*), n=3. (**F**) Volcano plot depicting changes in protein expression between integrated EF-pSer variant (TRC*, proD) strains and cells with OXB20-EF-pSer integration. (**G**) Volcano plot depicting changes in protein expression between integrated RS/EF strain and cells with no integrated OTS components (WT). Proteins above the curved lines represent significant expression changes; n = 3. (**H**) Schematic of tRNA cassette integration at yzgL and tRNA variations tested to improve tRNA orthogonality. (**I**) Mass spectrometry-based reporter for translation fidelity at TAG codons in integrated tRNA variant strains. Bar graph displaying values for Gly, Ser, and Thr misincorporation relative to pSer (represented by the non-phospho band on the PhosTag immunoblot), n=3. (**J**) Integrated tRNA variant strains were screened for their ability to facilitate pSer-mediated PPIs using a known pSer-dependent 14-3-3 PPI reporter. Unshifted Ser control population (grey) was compared to shifted productive PPI mediated by pSerOTS function (red or green).

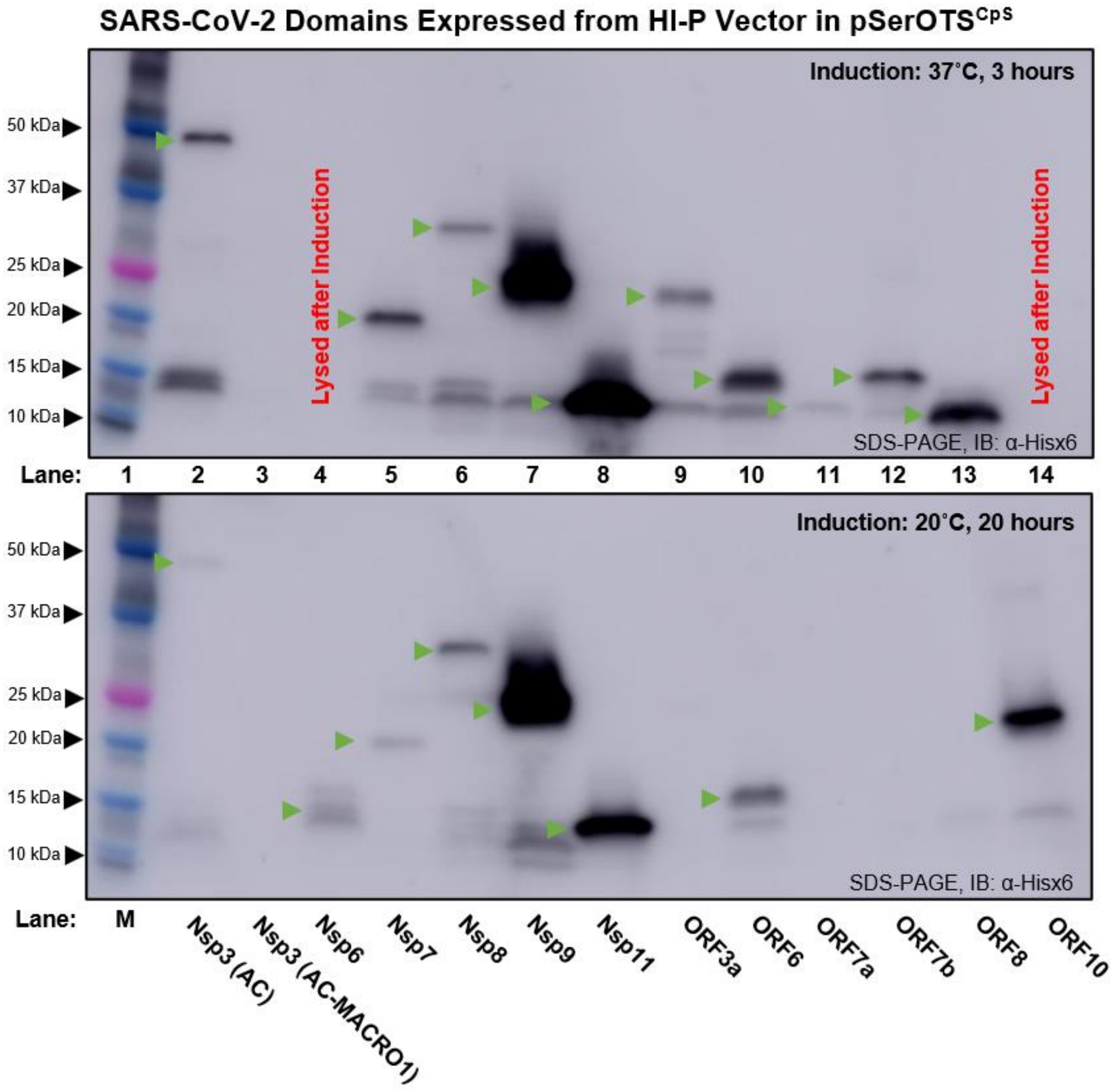


**Fig. S3. Expression of SARS-CoV-2 proteins from H-PIP platform.** The expression of SARS-CoV-2 proteins from H-PIP vectors was assessed by immunoblot against the 6xHis tagged split mCherry-viral fusion protein. Protein expression was carried out at 37 ºC for 3 hours (top) or 20 ºC for 20 hours (bottom) and an equal amount of OD normalized cell lysate was loaded in each lane.

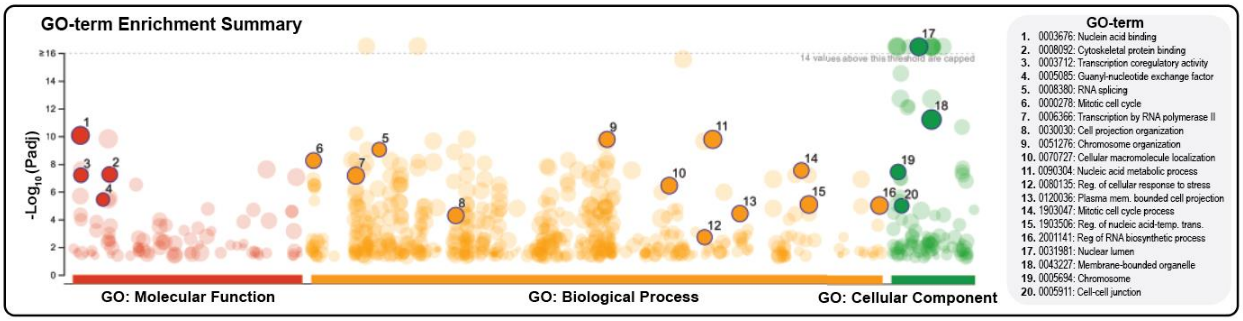


**Fig. S4. GO enrichment analysis of cumulative viral PPI network.** Proteins identified through H-PIP network analysis in Fig. 1A were used as input for GO enrichment analysis using g:Profiler. Nodes represent terms with adjusted p-values < 0.05, with node size reflecting the number proteins associated with an individual term related to molecular function (red), biological process (orange), or cellular component (green). Bold, highlighted GO-terms are listed.

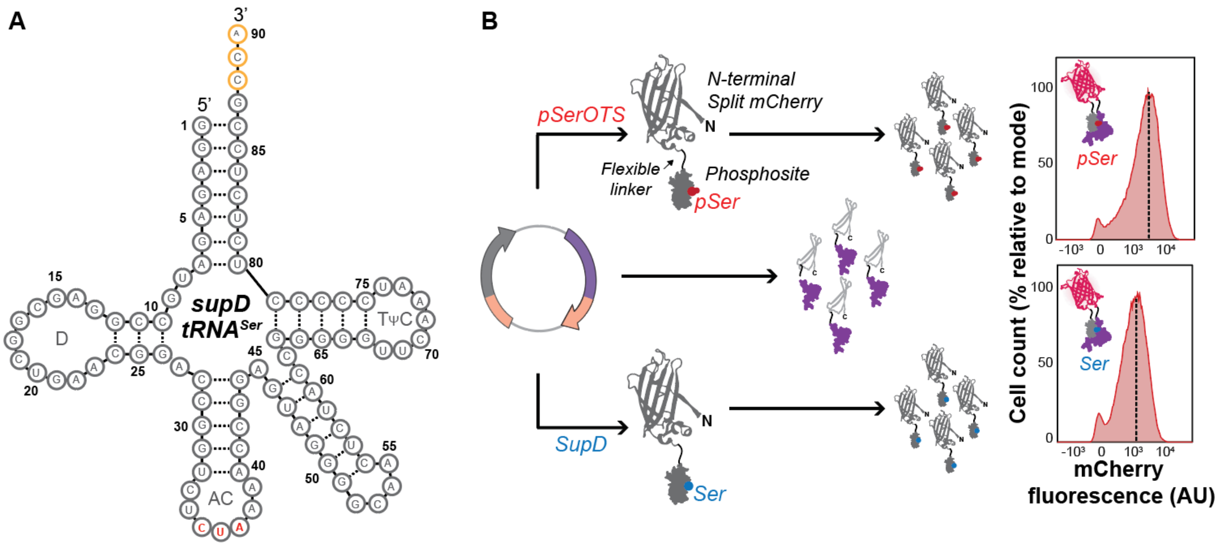


**Figure S5. Identification of phospho-specificity using supD suppressor tRNA.** (**A**) Amber suppressor tRNA (*supD*) is a naturally occurring suppressor tRNA derived from *E. coli* tRNA$^{Ser}$ that has a mutated anticodon which enables translation of UAG codons as Ser. (**B**) When used in place of pSerOTS, the phosphosite library can be expressed as a non-phosphorylated version of each "phosphosite." Comparison of PPIs identified in the Ser interactome to those obtained from the pSer interactome for each viral protein allows for the identification of pSer-specific PPIs.

**A** ORF7b: 1 - **MIELSLIDFYLCFLAFLLFLVLIMLIIFWFSLELQDHNETCHA** - 43

| Helix | Hydrophobicity | Hydrophobic Moment | Net Charge (z) | Polar Residues (n / %) | Nonpolar Residues (n / %) |
|---|---|---|---|---|---|
| 1 | 1.05093 | 0.08456 | -5 | 12 / 27.91% | 31 / 72.09% |

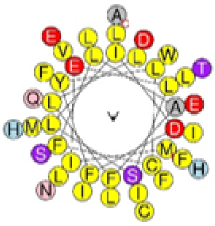

**B**

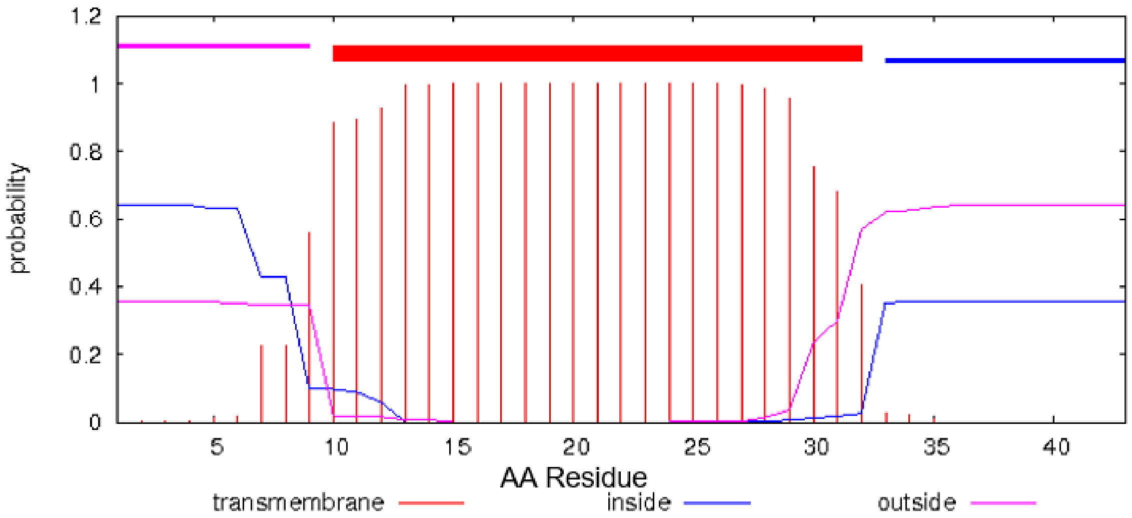


**Figure S6. Bioinformatic analysis of ORF7b biophysical properties.** (**A**) Biophysical properties for ORF7b were determined *in silico* from the primary amino acid sequence. Secondary structure prediction, charge state, and hydrophobicity were calculated to determine helicity and amphipathic potential. (**B**) Transmembrane potential, assessed using TMHMM, identified a single transmembrane helix with high probability spanning ~F9-L31.

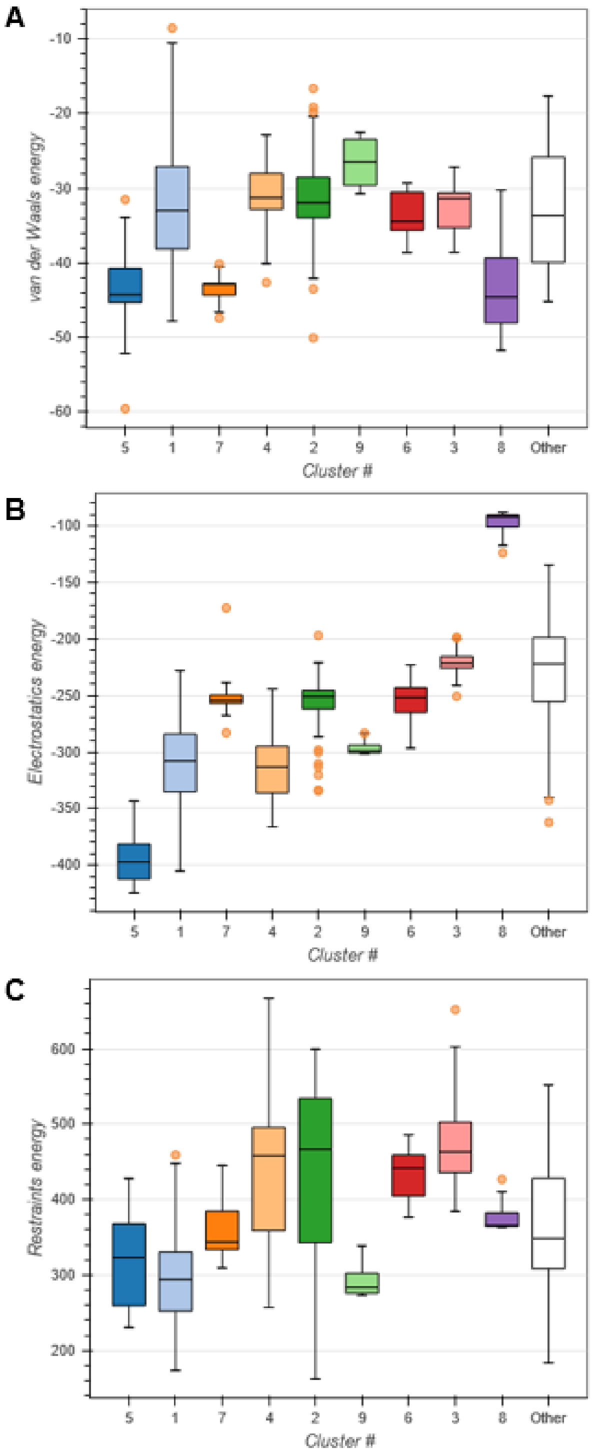


**Figure S7. Cluster analysis of ORF7b-Nucleocapsid docking models.** Iterative models of molecular interactions from HADDOCK molecular docking analysis were clustered based on (**A**) Van der Waals energy, (**B**) electrostatic energy, and (**C**) restraint energy and used to determine high-confidence docking models.

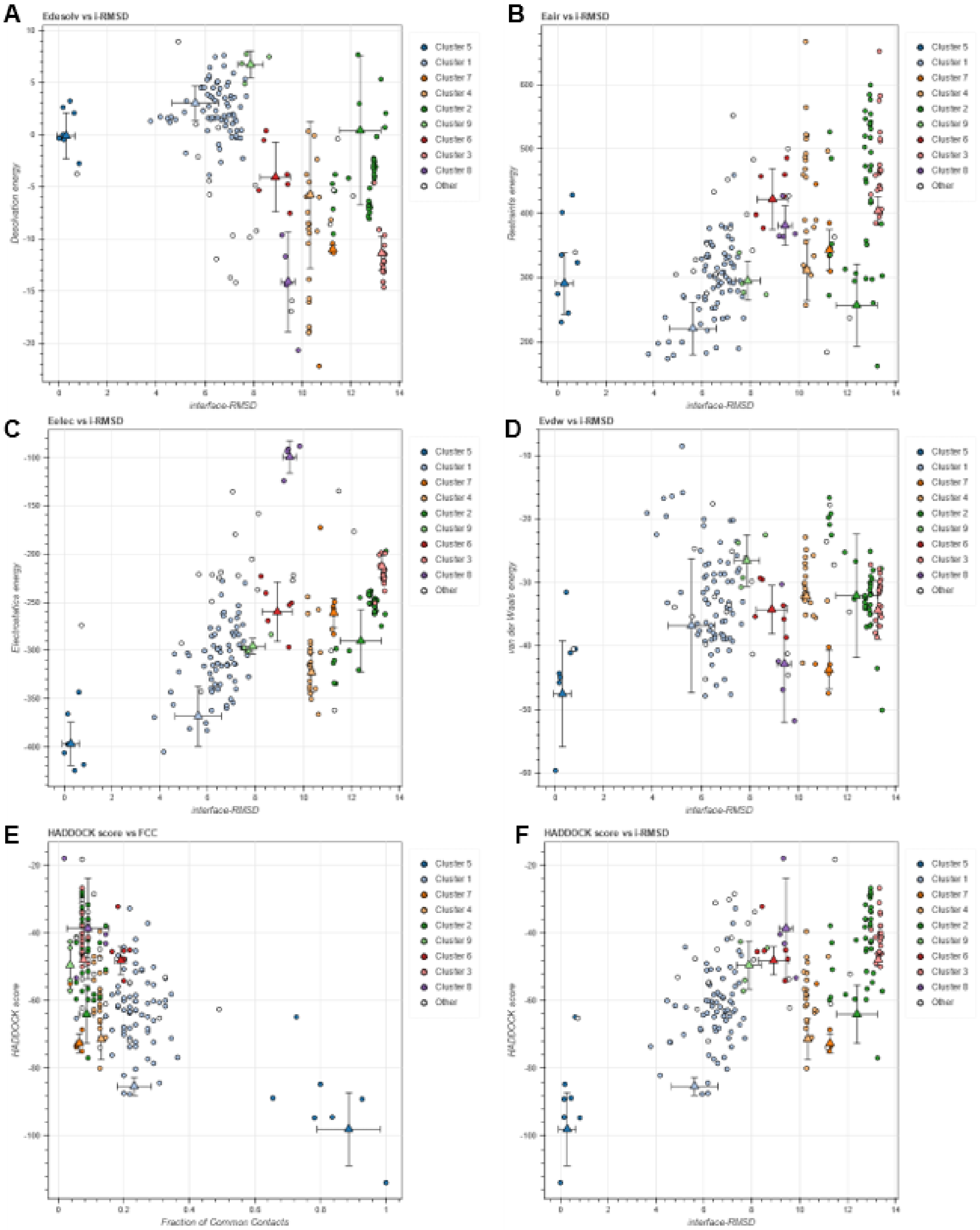


**Figure S8. Pairwise comparison of scoring from individual molecular docking iterations.** Docking model iterations were clustered by principal component analysis. (**A-F**) Pairwise comparison of computed biophysical interaction metrics from HADDOCK simulations.

| Strains | Name/Description | Growth Requirements | Reference |
|---|---|---|---|
| | rEcoli$^{XpS}$ (MG1655-C321 mutS+, λ-, Δ(ybhB-bioAB)::zeoR, ΔprfA, ΔserB) | No AB required (zeo resistant), requires d-biotin | (Mohler, 2021) |
| | rEcoli$^{CpS}$ (rEcoli$^{XpS}$, cybC::proD-pSerRS, glvC::TRC*-EF-pSer, yzgL::2xtRNA$^{pSer}$) | No AB required (zeo resistant), requires d-biotin | This Study |
| **Plasmids** | **Name/Description** | **Growth Requirements** | **Reference** |
| E29 | pSerOTSλ | Kanamycin, 37 ˚C | (Pirman, 2015) |
| G30 | supD tRNA$^{Ser}$ Suppressor | Kanamycin, 37 ˚C | (Pirman, 2015) |
| Q81 | glnS*-pSerRS FRT-KAN-FRT | Kanamycin, 37 ˚C | (Mohler, 2021) |
| U66 | TRC*-pSerRS FRT-KAN-FRT | Kanamycin, 37 ˚C | This Study |
| U67 | proD-pSerRS FRT-KAN-FRT | Kanamycin, 37 ˚C | This Study |
| U68 | OXB20-pSerRS FRT-KAN-FRT | Kanamycin, 37 ˚C | This Study |
| V68 | TRC*-EF-pSer21 FRT-KAN-FRT | Kanamycin, 37 ˚C | This Study |
| V66 | proD-EF-pSer21 FRT-KAN-FRT | Kanamycin, 37 ˚C | This Study |
| V67 | OXB20-EF-pSer21 FRT-KAN-FRT | Kanamycin, 37 ˚C | This Study |
| W54 | RK6, 2x tRNApSer G2-C71 Cassette FRT-KAN-FRT | Kanamycin, 37 ˚C | This Study |
| X60 | RK6, 2x tRNApSer U2-A71 Cassette FRT-KAN-FRT | Kanamycin, 37 ˚C | This Study |
| O23 | (pCP20) Rep101(Ts), FLP recombinase | Kanamycin, 30 ˚C | (Wanner, 2000) |
| O34 | (pKD46) Rep101(Ts), bet-gam-exo lambda phage | Kanamycin, 30 ˚C | (Wanner, 2000) |
| O32 | (pKD4) RK6, FRT-KAN-FRT | Kanamycin, 37 ˚C | (Wanner, 2000) |
| | *Recombinant Reporter Expression* | | |
| N54 | HI-P split mCherry reporter with 14-3-3β and PSP3-7, IPTG/Ara Inducible, p15a | Ampicillin, 37 ˚C | (Barber, 2018) |
| E52 | TAG-MS-READ mass spectrometry reporter, aTc Inducible, p15a | Ampicillin, 37 ˚C | (Mohler, 2017) |

**Table S2. Plasmids and Strains used in this study.**

**Data S1. (separate file)**

Supplementary data file containing the processed NGS data for each SARS-CoV-2 protein analyzed by H-PIP.

**Data S2. (separate file)**

Supplementary data file for GO enrichment analysis using gProfiler presented in Figure S3 based on PPI network in Figure 1A.
